# How Generative Models Approach Molecular Conformational Sampling

**DOI:** 10.64898/2026.04.10.717851

**Authors:** B E Nagesh, Jagannath Mondal

## Abstract

Characterising equilibrium conformational ensembles with deep generative models requires assessing not only whether a model reproduces the target distribution, but also the mechanism of *how* it arrives here. Here we examine two distinct routes to generative conformational sampling— stochastic relaxation and deterministic transport—through a study of denoising diffusion probabilistic models (DDPM) and rectified-flow (RF) models across molecular systems of increasing complexity. Using systems of increasing complexity, including a multimodal two-dimensional potential, the folded mini-protein Trp-cage, and a high-dimensional dihedral subspace of the intrinsically disordered protein *α*-synuclein, we show that the key distinction between these paradigms lies not only in endpoint fidelity but in how distributional error is resolved during sampling. Diffusion models converge via pronounced late-stage stochastic relaxation and exhibits robust recovery of configurational breadth across neural architectures. Rectified flow approaches the target more gradually through deterministic transport and therefore depends much more strongly on architectural expressivity, particularly in heterogeneous high-dimensional landscapes. Analyses of entropy and moment evolution further show that diffusion more reliably restores both ensemble location and fluctuation structure, whereas RF requires Transformer-level feature mixing to represent the transport geometry accurately. These results establish convergence mechanism as a key design principle for generative sampling.

## I. INTRODUCTION

Molecular dynamics (MD) simulations provide atomistic access to conformational ensembles, but their central limitation is sampling efficiency: free-energy landscapes with multiple metastable basins, broad disordered regions, or slow collective rearrangements render converged equilibrium sampling prohibitively expensive, particularly for intrinsically disordered proteins and other systems with strongly coupled degrees of freedom [1–4]. Deep generative models offer a complementary strategy: rather than propagating trajectories, they aim to learn the equilibrium distribution from finite data and generate new configurations directly from it. In this view, generative models are learned samplers of conformational space, capable of augmenting conventional MD while preserving the statistical structure of the underlying ensemble.

Two prominent continuous-time generative paradigms have emerged: diffusion models [5– 10] and flow-based models [11–15]. Denoising diffusion probabilistic models (DDPM) [5, 8] generate samples by reversing a stochastic noising process, combining a learned denoising field with stochastic relaxation during sampling. Rectified flow (RF) [13], the flow-based representative studied here, instead learns a deterministic velocity field that transports samples from a Gaussian base to the data distribution via straight-line trajectories in state space. Both paradigms can, in principle, represent complex probability densities—but they do so through fundamentally different dynamical mechanisms, and this difference has consequences that extend well beyond final sample quality.

The question we ask here is not which model is better, but rather: *how does each model arrive at its generated ensemble, and what does that route reveal about the underlying dynamics?* This distinction matters because the route to the target distribution—the full trajectory of convergence—is directly tied to robustness, sensitivity to approximation error, and the architectural capacity required for faithful sampling. A model that achieves low KL divergence through stochastic relaxation is mechanistically different from one that achieves the same endpoint through accurate deterministic transport, even if their final samples are superficially indistinguishable.

A second, equally important axis of the present study concerns neural architecture [16–20]. Conformational spaces are highly structured and correlated, and the success of a generative model cannot be separated from the expressive capacity of the network representing it. Multilayer perceptrons (MLPs) [18] provide a minimal baseline; residual MLPs improve optimization stability and enable deeper representations; Transformer architectures [20] model long-range dependencies through self-attention. Architecture is therefore not a secondary implementation detail but an integral part of the scientific question: how does generative dynamics interact with representational capacity to shape the *entire* convergence process?

Despite rapid progress in molecular generative modeling—including Boltzmann generators [21], diffusion-based protein sampling [22, 23], and energy-based generative approaches [24]—the mechanistic basis for convergence has remained largely unexplored. Prior comparative studies have examined diffusion, flow-matching, and related approaches primarily at the level of hyperparameter sensitivity and benchmark accuracy [25]. In particular, how stochastic versus deterministic dynamics interact with architecture to produce their respective sampling trajectories has not been systematically examined in molecular settings.

Here, we address this gap by analyzing not only endpoint distributions but the full time evolution of KL divergence, distributional moments, and entropy, comparing denoising diffusion probabilistic models (DDPM) [8] and rectified-flow (RF) models [13] across three systems of increasing complexity: a two-dimensional three-well potential, the folded mini-protein Trp-cage in a 38-dimensional dihedral space, and a 60-dimensional dihedral subspace of the intrinsically disordered protein *α*-synuclein (Fig. 1). For each paradigm, we consider three neural architectures of increasing representational capacity (Fig. 2): a standard MLP, a residual MLP (MLP-RC), and a Transformer. By examining *how* diffusion and RF approach the target distribution rather than merely *where* they end up, we uncover a mechanistic and architectural picture that is both more informative and more actionable than endpoint comparisons alone. We show that diffusion achieves convergence through late-stage stochastic relaxation—a dynamical mechanism with an intrinsic dissipative character—whereas RF transports probability mass gradually and deterministically, with convergence tied directly to the fidelity of the learned velocity field. This mechanistic distinction explains, in a unified way, why diffusion is comparatively insensitive to neural architecture while RF becomes progressively more dependent on model expressivity as the conformational landscape grows broader and more correlated.

**FIG. 1.**
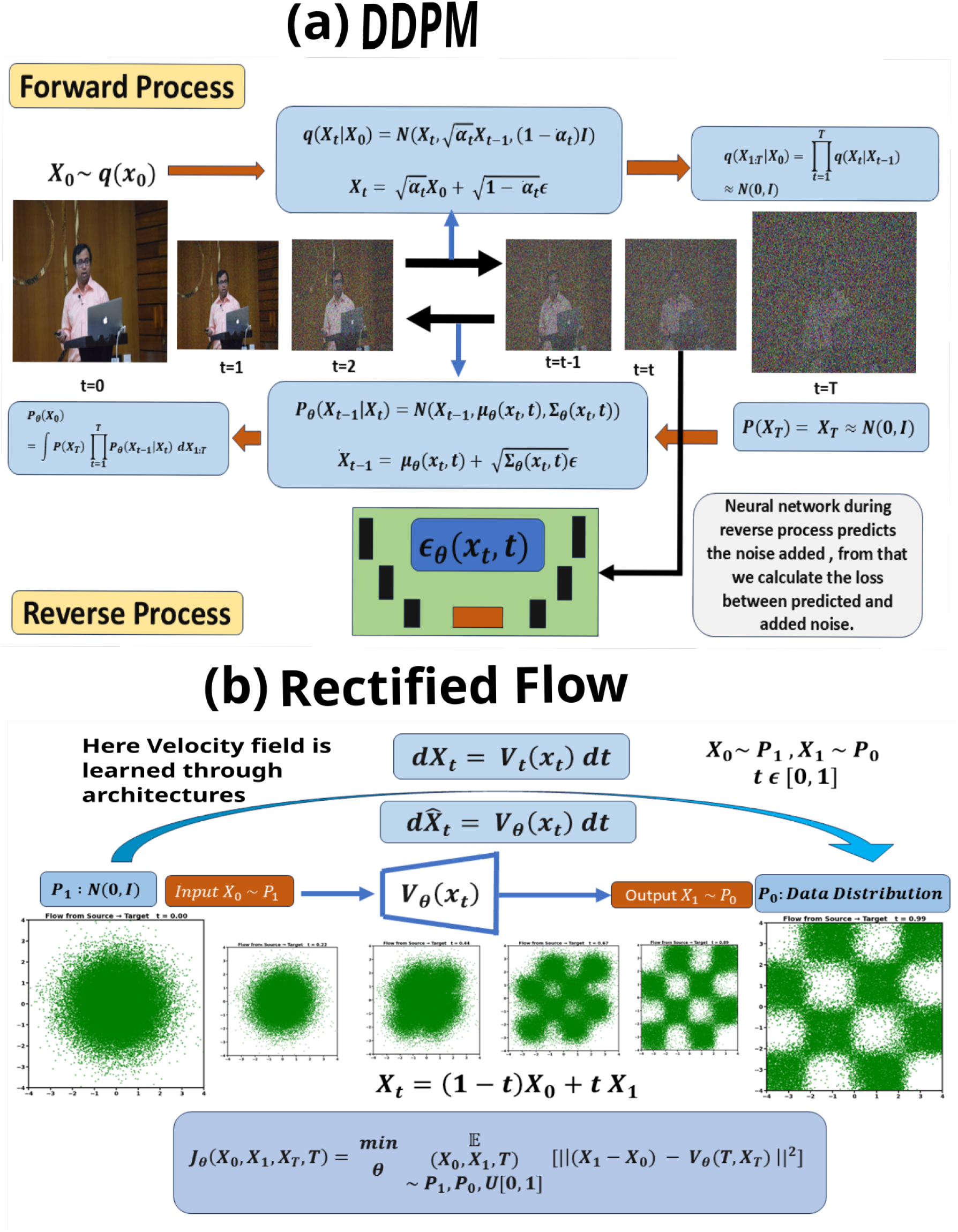
Schematic representation of the two generative paradigms. (a) In denoising diffusion probabilistic models (DDPM), a fixed forward noising process is paired with a learned reverse denoising process. (The model was trained on image of one of the authors of this manuscript with full consent). (b) In rectified flow (RF), the model learns a deterministic velocity field that continuously transports samples from a base distribution to the data distribution. As shown in the results below, these distinct mathematical structures are directly reflected in the observed sampling dynamics and KL-divergence profiles.

**FIG. 2.**
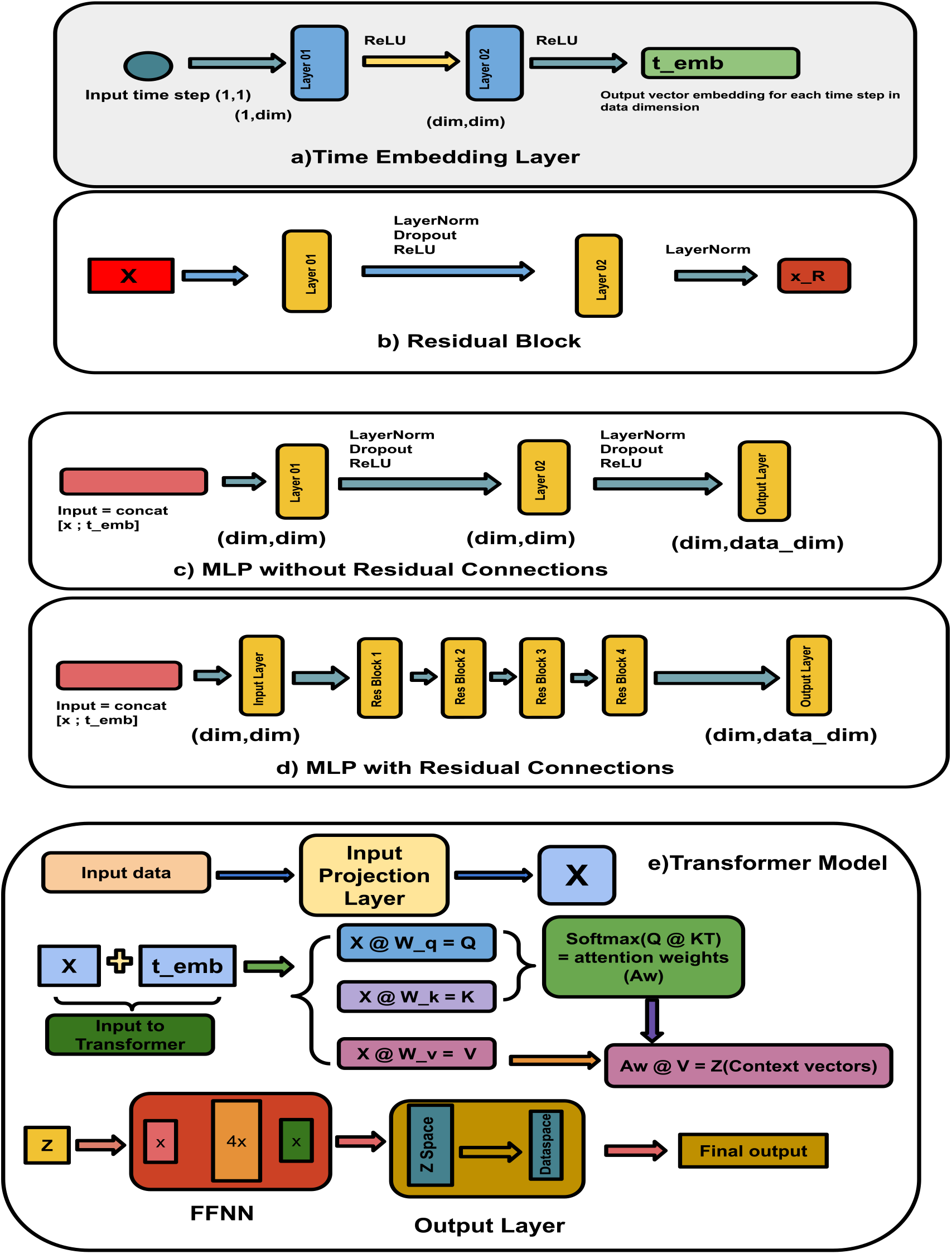
Neural architectures used for diffusion and rectified flow. (a) Time embedding used to encode the sampling coordinate. (b) Residual block used in the MLP-RC architecture. (c) Baseline MLP architecture. (d) Residual MLP architecture composed of stacked residual blocks. (e) Transformer-based architecture with self-attention and residual pathways. The progression from MLP to MLP-RC to Transformer defines a controlled hierarchy of increasing representational capacity. Under the mechanistic framework of this work, this hierarchy operationalizes the ex-pressivity requirement predicted by the generative dynamics: diffusion’s stochastic relaxation is where *d* is the data dimensionality. Depending on the generative paradigm, *f*_*θ*_ represents either the noise-prediction field *ϵ*_*θ*_(*x, t*) in DDPM or the velocity field *v*_*θ*_(*x, t*) in rectified flow.

## II. THEORY AND MODEL FORMULATION

### A. Density evolution as a unifying perspective

Let *x* ∈ 𝒳 denote a molecular conformation represented in a chosen coordinate system. In this work, 𝒳= ℝ ^2^ for the three-well potential and 𝒳= ℝ^*D*^ for the protein dihedral representations. The target distribution is denoted by *p*_data_(*x*), and the base distribution by *p*_base_(*z*), which is taken to be a standard Gaussian.

A useful common starting point for both diffusion and flow-based generative models is the time evolution of a probability density *p*_*t*_(*x*). Consider the stochastic differential equation

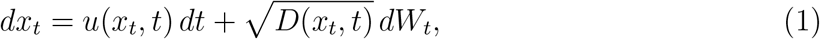

where *u*(*x, t*) is a drift field, *D*(*x, t*) is a diffusion coefficient, and *W*_*t*_ denotes a Wiener process. The corresponding density evolves according to the Fokker–Planck equation

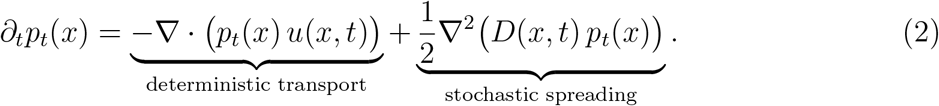

Equation (2) makes the key mechanistic distinction explicit. The Laplacian term introduces entropy production into the density evolution: it is strictly non-negative in its contribution to relative entropy decay and provides an intrinsic dissipative tendency that drives *p*_*t*_ toward equilibrium regardless of the accuracy of the drift *u*. Diffusion models exploit both terms. Rectified flow corresponds to the deterministic limit *D*(*x, t*) = 0, in which the Laplacian term vanishes entirely and density evolution is governed solely by the continuity equation. This is not merely a formal difference: it means that diffusion dynamics possess a built-in self-correcting mechanism that deterministic transport lacks. As we show in the results, this distinction is directly visible in the KL-divergence trajectory during sampling—a pronounced late-stage relaxation in diffusion versus a gradual transport-driven convergence in RF—and it determines how strongly each paradigm depends on neural architectural expressivity.

### B. Score-based diffusion models

In the variance-preserving formulation, the forward diffusion process is defined by

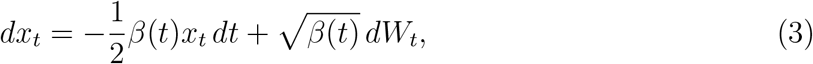

where *β*(*t*) is a prescribed noise schedule. This process progressively transforms the data distribution into a simple Gaussian reference distribution, establishing the endpoint from which the learned reverse process must recover the data.

(Throughout the article, we use the term “diffusion” and “DDPM” interchangeably. On the other hand we use the term ‘Flow’, “RF” or “Rectified flow’ interchangably,) In the discrete-time DDPM formulation [8], the forward transition is

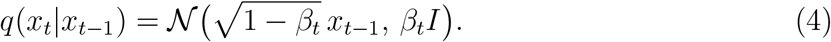

Defining

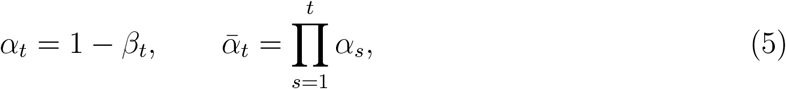

one obtains the marginal form

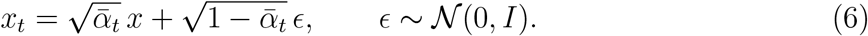

The reverse process transforms noise back into data and, in continuous time, may be written as

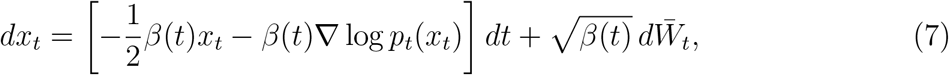

where the score function ∇ log *p*_*t*_(*x*) is approximated using a neural network *s*_*θ*_(*x, t*). In practice, DDPM is commonly parameterized through noise prediction,

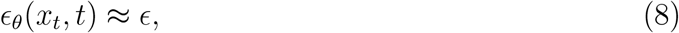

which is equivalent to score prediction through

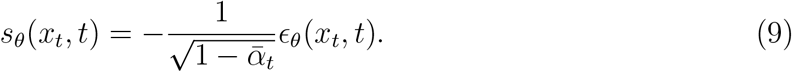

The corresponding training objective is

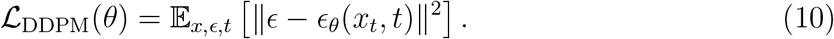

The mechanistic consequence for sampling is the following. Diffusion does not learn transport directly; it learns a denoising field embedded within a stochastic reverse process whose dynamics retain the Laplacian entropy-producing term. Sampling therefore combines learned drift with stochastic relaxation. Because the dynamics intrinsically contract relative entropy—independently of how well the neural network approximates the score—the reverse process can redistribute probability mass into the correct basins even when the learned field is imperfect. This is the origin of diffusion’s architectural robustness: the network does not need to represent the full global transport map; it provides local denoising directions within a process that self-corrects.

### C. Rectified-flow models

Rectified flow (RF) [13] defines a deterministic trajectory in configuration space,

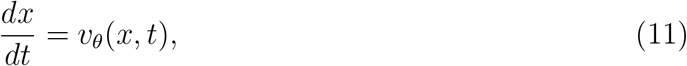

with density evolution governed by the continuity equation

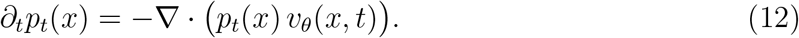

A linear interpolation between a base sample *z* ∼ *p*_base_ and a data sample *x* ∼ *p*_data_ is introduced as

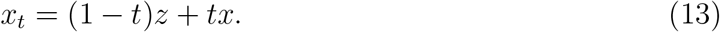

The corresponding target velocity field is

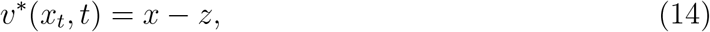

and the neural network is trained to approximate the conditional expectation of this velocity,

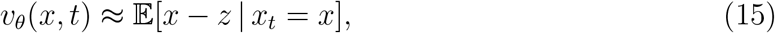

using the objective

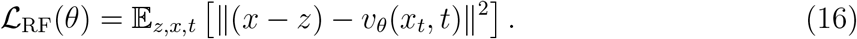

The mechanistic consequence is the direct counterpart to diffusion. Because Eq. (12) contains no Laplacian term, the time derivative of the KL divergence between the generated density and the target has no guaranteed sign:

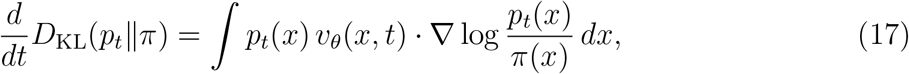

whose sign depends entirely on the learned velocity field. There is no intrinsic dissipative mechanism; convergence must be achieved entirely through the accuracy of *v*_*θ*_. Any error in the velocity field propagates without correction, and the generated distribution at *t* = 1 directly reflects the accumulated approximation error. This places a fundamentally greater burden on architectural expressivity than diffusion: the network must represent a coherent global transport field, not merely local denoising directions. As a consequence, the quality of RF sampling is architecture-dependent in a way that diffusion’s is not, and this dependence grows more severe as the target ensemble becomes higher-dimensional, more correlated, and more heterogeneous.

We refer to the Supplementary Information (Sections I and II) for a full derivation of diffusion and flow models from the Fokker–Planck framework, including the unified derivation of KL-divergence convergence behaviour for both paradigms.

### D. Architectures and molecular systems

The theoretical distinction above—stochastic relaxation in diffusion versus architecture-dependent deterministic transport in RF—immediately implies a prediction about neural architecture: the expressivity requirement for RF is set by the dynamics themselves, not by implementation convenience. To test this prediction systematically, we consider three neural architectures of increasing representational capacity: a multilayer perceptron (MLP) [18], a residual multilayer perceptron (MLP-RC), and a Transformer [20] (Fig. 2).

The MLP provides the simplest baseline for learning time-conditioned score or velocity fields, with limited ability to represent cross-dimensional correlations. The MLP-RC augments this with residual connections that improve optimization stability and enable deeper representations while preserving the feedforward character. The Transformer employs self-attention-based feature mixing and residual pathways, making it structurally suited to represent correlated structure across dimensions—the kind of global feature mixing that deterministic transport requires when the target ensemble is correlated. Under the new narrative, the three architectures are not merely a performance ladder; they operationalize the expres-sivity axis along which the dynamical distinction between diffusion and RF is expected to manifest.

All architectures are explicitly conditioned on time and output a vector field in the same dimensionality as the input coordinates. This time-dependent function corresponds to *ϵ*_*θ*_(*x, t*) for DDPM and to *v*_*θ*_(*x, t*) for rectified flow. A full layer-by-layer specification of each architecture is provided in the Supplementary Information.

We evaluate these models on three systems of increasing complexity. First, we consider a particle evolving in a two-dimensional three-well potential, which provides a low-dimensional benchmark where the full free-energy surface can be visualized directly and KL divergence can be tracked throughout sampling. The training data for this system are generated from molecular dynamics simulations performed in our group [26, 27]. Second, we study the folded mini-protein Trp-cage, represented in a 38-dimensional backbone dihedral space consisting of 19 (*ϕ, ψ*) pairs. For this system, we use long molecular dynamics trajectories from D. E. Shaw Research (DESRES), spanning 100 *µ*s and generated using the a99SB-disp force field on Anton hardware; detailed simulation protocols are available in prior work [28, 29]. Finally, we examine a 60-dimensional dihedral subspace of the intrinsically disordered protein *α*-synuclein, corresponding to 30 (*ϕ, ψ*) pairs. We again utilize DESRES trajectories, including an extended 73 *µ*s simulation that mitigates periodic imaging artifacts present in earlier datasets [28, 30].

Together, these three systems define a controlled progression from low-dimensional multimodality to correlated folded-protein ensembles and finally to a broad intrinsically disordered ensemble. The hierarchy is designed specifically to ask how the mechanistic distinction between stochastic relaxation and deterministic transport scales with dimensionality, conformational heterogeneity, and architectural capacity—with the expectation, grounded in the theory above, that the gap between diffusion and RF will widen as the transport problem becomes more global.

### E. Model implementation and training protocol

All models parameterize a time-dependent function

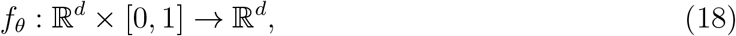

All architectures use the same hidden dimensionality and are trained under a matched optimization protocol, ensuring that any observed differences in sampling fidelity and convergence dynamics can be attributed to the interaction between generative mechanism and representational capacity rather than to unrelated optimization choices. In the present implementation, we use a hidden dimension of 256, six Transformer layers with eight attention heads, a dropout rate of 0.2, a batch size of 4096, and a learning rate of 5 × 10^−5^. Models are trained for 50,000 epochs using the AdamW optimizer [31, 32] with weight decay 10^−4^ and are implemented in PyTorch [33].

## III. RESULTS

Our analysis focuses on the approach to the target equilibrium distribution rather than endpoint agreement alone. To this end, we use free-energy reconstruction, KL divergence, entropy difference, and moment evolution as complementary probes of how each generative paradigm redistributes probability mass during sampling. Across all systems examined, the central observation is that diffusion and rectified flow do not merely differ in final fidelity; they differ in the route by which that fidelity is achieved. Diffusion resolves error through stochastic relaxation, with distributional mismatch often collapsing sharply at later stages of generation, whereas rectified flow converges more gradually through deterministic transport and succeeds only to the extent that the learned velocity field accurately captures the geometry of the target ensemble. The results below show that this distinction governs both robustness to architecture and performance across increasingly heterogeneous molecular landscapes.

Before turning to the system-specific analysis, it is useful to recall what is being tested by the three architectures in Fig. 2. The baseline MLP provides the simplest parameterization of the time-dependent field learned by the model and therefore serves as a reference for low-capacity generative modeling. The residual MLP improves optimization stability and representational depth while preserving the same basic feedforward character. The Transformer changes the modeling assumption more substantially: through self-attention it enables each coordinate to condition on all others, which is especially relevant when the target distribution contains strongly coupled degrees of freedom. The role of architecture in the following sections should therefore be interpreted not as a purely technical choice, but as a test of how much expressive power is required for each generative mechanism to approach the target distribution reliably.

### A. A low-dimensional landscape reveals distinct requirements for approaching equilibrium

We begin with a two-dimensional three-well potential, which provides a transparent setting in which the full free-energy surface (FES) can be visualized directly without dimensional reduction. The reference dataset consists of *N* = 5 × 10^5^ samples, yielding a well-converged ground-truth FES. To create a deliberately data-limited regime, only 10% of this dataset is used for training. This system serves as the clearest window into the mechanistic distinction between the two paradigms: because the full distribution is accessible, we can track not only endpoint quality but the complete convergence trajectory.

Both diffusion and rectified-flow (RF) models are trained using the same three architectures— MLP, MLP-RC, and Transformer—and evaluated according to their ability to reconstruct the reference distribution. The training subset already reveals incomplete coverage of the underlying landscape, making this a stringent test of generative recovery.

Figures 3(b)–(d) show the FES generated by diffusion models across architectures. All three architectures recover the global basin topology with good qualitative agreement. To quantify this agreement, we compute the Kullback–Leibler (KL) divergence [34],

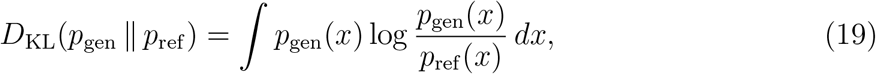

where *p*_ref_ (*x*) denotes the ground-truth distribution and *p*_gen_(*x*) denotes the generated one. Across architectures, diffusion achieves uniformly low KL divergence (Fig. 3f), indicating accurate recovery of probability mass even with comparatively simple models. The residual architecture performs most consistently, but the overall dependence on architecture remains weak. This is not coincidental: as we show below, the stochastic reverse process provides an intrinsic relaxation tendency that partially compensates for imperfections in the learned denoising field, making the outcome comparatively insensitive to model capacity.

**FIG. 3.**
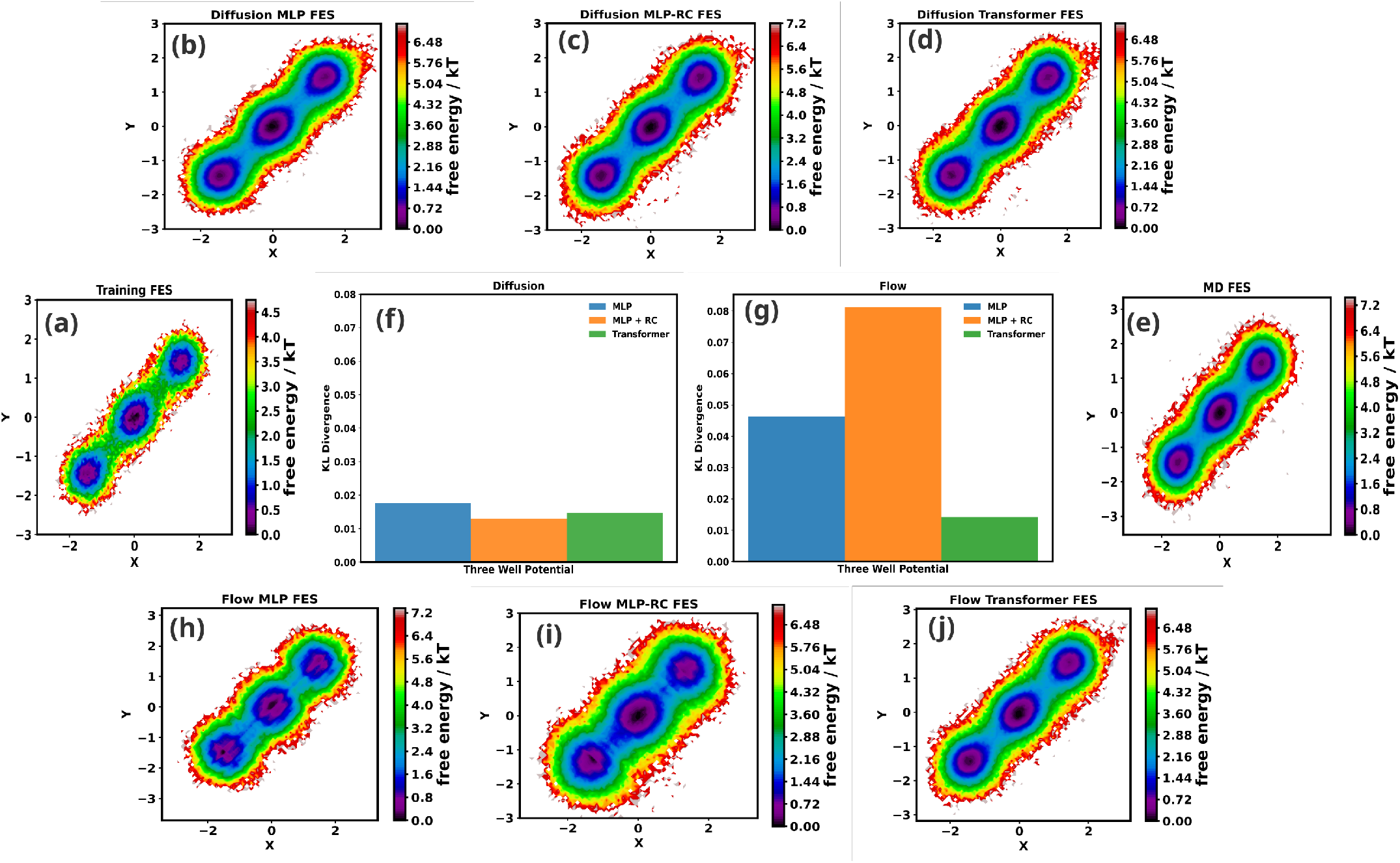
Sampling of the three-well potential with diffusion and rectified-flow models. Panels (a) and (e) show the training subset and reference free-energy surface, respectively. Panels (b)–(d) show the generated free-energy surfaces for MLP, MLP-RC, and Transformer architectures using diffusion. All three architectures recover the global basin topology, with the residual architecture giving the most consistent quantitative agreement, reflecting the robustness that stochastic relaxation confers even on moderate-capacity networks. Panel (f) shows the KL divergence between the reference and diffusion-generated distributions. Panels (h)–(j) show the generated surfaces for the same architectures using rectified flow. In contrast to diffusion, RF depends strongly on architecture because deterministic transport must represent the full basin geometry without a self-correcting mechanism; only the Transformer does so with high fidelity. Panel (g) shows the corresponding KL divergence for RF.

In contrast, RF exhibits a pronounced sensitivity to architectural capacity. Figures 3(h)–(j) show that while the Transformer reproduces the basin structure with high fidelity, the MLP and MLP-RC models fail to resolve the wells sharply and misallocate probability across basins. This difference is reflected in the KL divergence (Fig. 3g), where RF with Transformer substantially outperforms the MLP-based variants. Because RF sampling is entirely deterministic, there is no intrinsic mechanism to correct errors in the learned velocity field: the architecture must represent the global transport geometry accurately, and when it does not, the resulting distribution reflects those errors directly.

Even in this low-dimensional system, therefore, the two paradigms display qualitatively different architecture dependence—a distinction rooted in their underlying dynamics rather than in any surface-level property of the models.

### B. Trp-cage: the mechanistic gap widens in correlated protein conformational space

We next test whether the architecture robustness of diffusion and the capacity sensitivity of RF persist and deepen as we move to a realistic protein system. Trp-cage provides an informative intermediate regime: the conformational space is substantially higher-dimensional and more correlated than the three-well potential, yet still sufficiently structured to permit detailed analysis of pairwise marginals, KL divergence, and entropy preservation. Crucially, this system also allows us to examine how each paradigm approaches the target distribution during sampling via moment evolution, providing the first mechanistic window into convergence dynamics in a high-dimensional setting.

The system is represented in a 38-dimensional backbone dihedral space comprising 19 (*ϕ, ψ*) pairs,

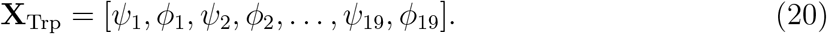

We use long molecular dynamics trajectories (100 *µ*s) from D. E. Shaw Research as the reference distribution, while restricting training to 10% of the available data to create a data-limited regime.

Figure 4(d,e) compares the FES obtained from the training subset and from the full trajectory, highlighting the incomplete coverage of conformational space in the training data. Figures 4(a)–(c) show representative (*ϕ, ψ*) marginal distributions generated by diffusion models across architectures. Diffusion reproduces the dominant conformational features with good fidelity across all three architectures. The MLP-RC model provides the most consistent agreement across residues, while the plain MLP exhibits minor smoothing and the Transformer does not yield a systematic improvement over MLP-RC—a pattern consis-tent with the stochastic relaxation mechanism providing a floor of robustness that makes additional architectural capacity less decisive. Additional representative pairs are provided in the Supporting Information (Figs. S1–S3), where the same trend persists across the full set of residues.

**FIG. 4.**
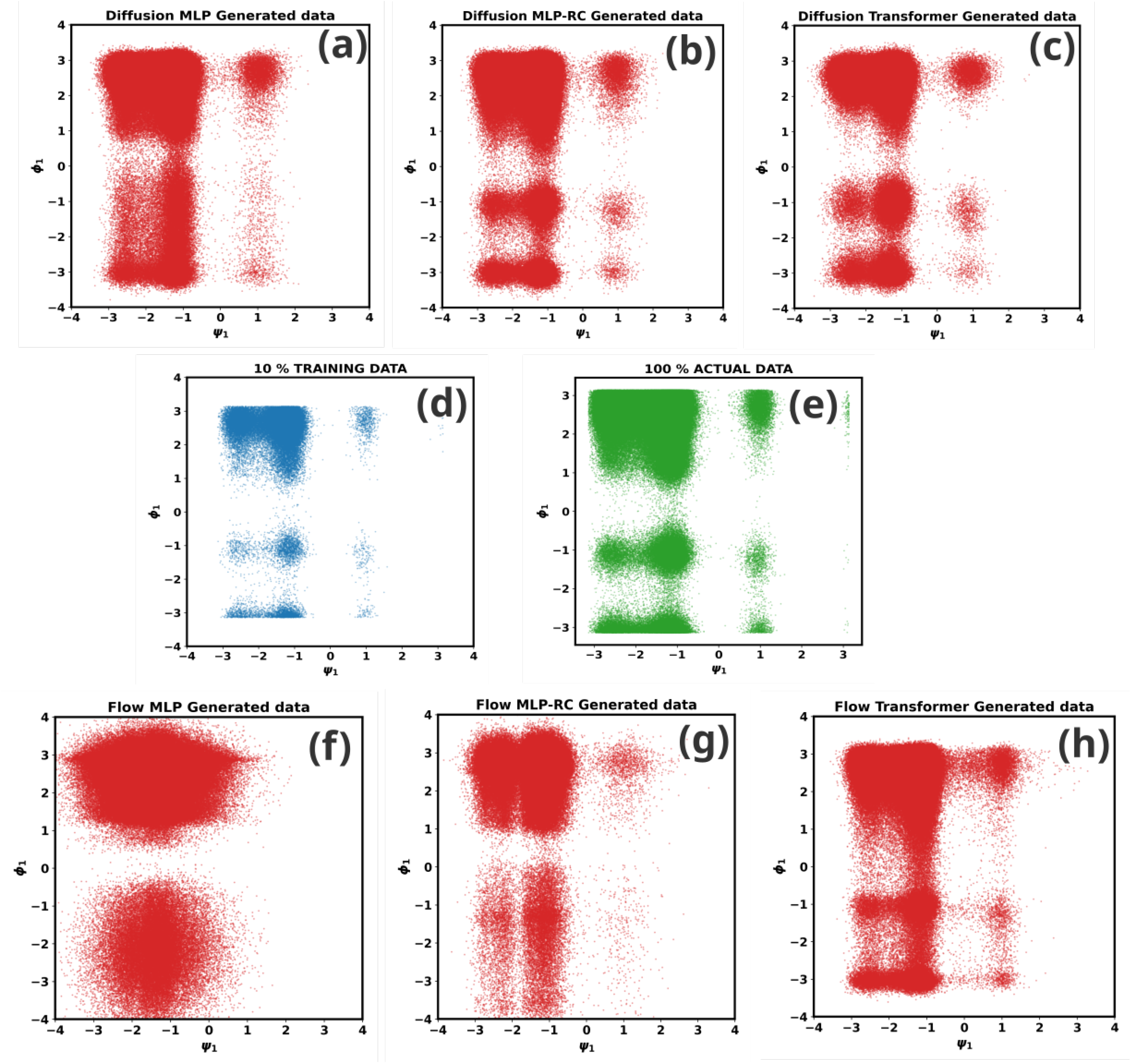
Representative (*ψ, ϕ*) marginals for Trp-cage. Panels (d) and (e) show the training subset and the full reference ensemble. Panels (a)–(c) correspond to diffusion models and panels (f)–(h) to rectified-flow models. Diffusion reproduces the marginal structures with comparatively weak dependence on architecture—a signature of stochastic relaxation compensating for limited model capacity—whereas RF displays a pronounced sensitivity to representational capacity and requires Transformer-level expressivity to recover the target pairwise structure reliably, because deterministic transport cannot self-correct errors in the learned velocity field.

By contrast, RF exhibits a substantially stronger dependence on architectural capacity (Fig. 4f–h). The MLP and MLP-RC models fail to reproduce sharp features in the marginal distributions and show clear distortions in probability density. Only the Transformer architecture restores the reference distributions with high fidelity. In the folded protein setting, the representational bottleneck of RF becomes visible already at intermediate dimensionality, consistent with the need for the learned velocity field to model cross-dimensional correlations coherently—a requirement that MLP-based architectures cannot fully satisfy.

These observations are quantified through KL divergence across all 19 dihedral pairs (Fig. 5). Diffusion remains in a comparatively low-KL regime across architectures, with MLP-RC performing most reliably. In contrast, RF shows substantially larger KL divergence for MLP-based models, while the Transformer yields a marked reduction across nearly all residues. This result highlights a clear expressivity bottleneck in RF that does not arise to the same degree in diffusion—and its origin is dynamical: without an intrinsic entropy production term, every error in the RF velocity field propagates uncorrected to the final distribution.

**FIG. 5.**
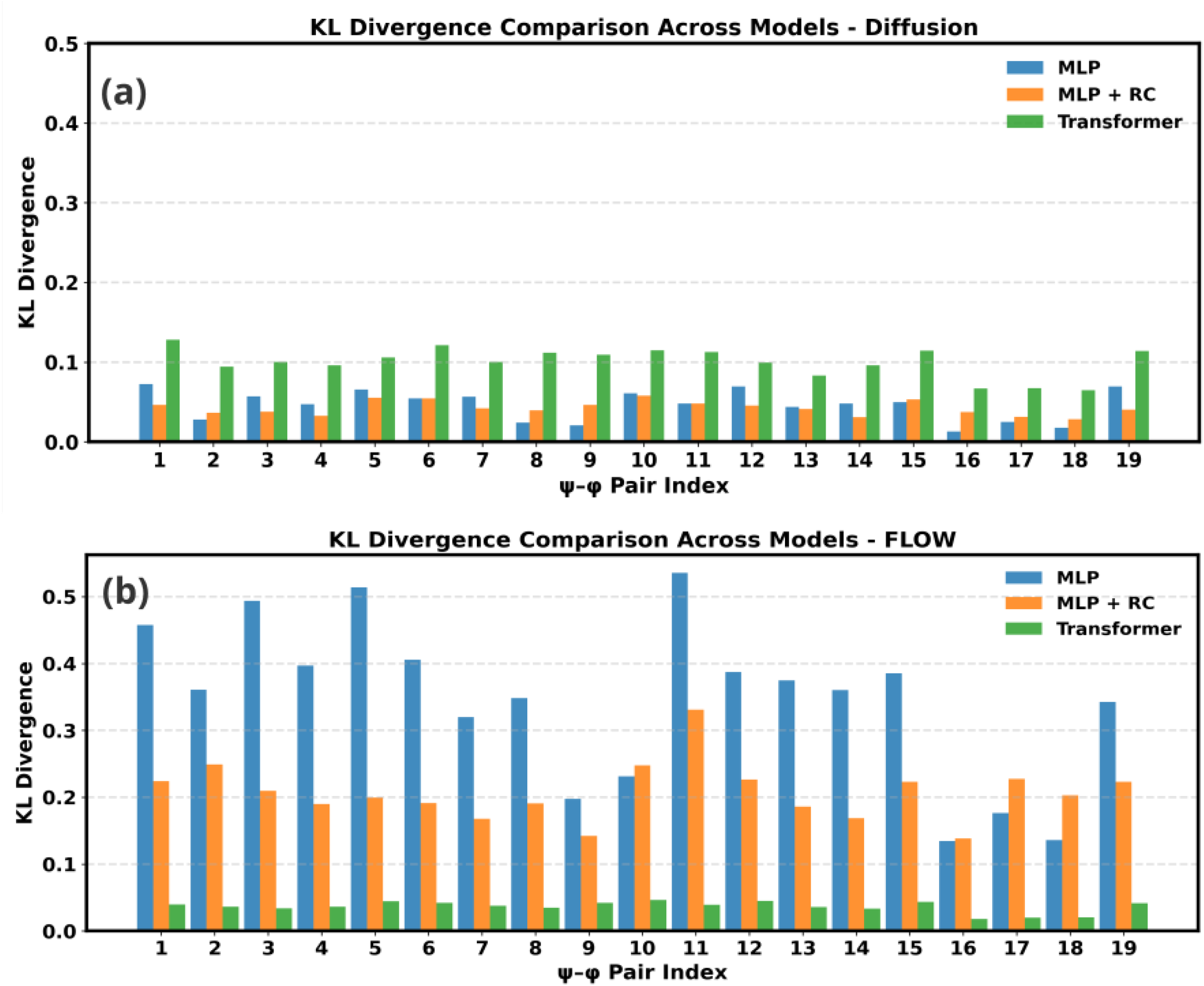
KL divergence across Trp-cage dihedral pairs. Panel (a) shows the diffusion results and panel (b) the rectified-flow results. Lower KL indicates better agreement between generated and reference pair distributions. Diffusion remains in a comparatively low-KL regime across architectures because stochastic relaxation provides a built-in correction mechanism that tolerates imperfect denoising fields. RF exhibits a pronounced capacity bottleneck—large KL for MLP-based models and a sharp reduction only with the Transformer—because deterministic transport requires the velocity field to represent the full distribution geometry accurately, with no self-correcting mechanism for approximation errors.

We also perform an error analysis by drawing ten independent samples for both diffusion and RF and computing the KL divergence of each against the reference distribution. The mean KL behaviour is shown in Fig. S4; the mean does not deviate substantially from individual KL values, confirming sampling consistency with the respective learned statistics. In Fig. S5, we increased the training data to 30% and observed that no marked improvement results, indicating that 10% already provides sufficient statistics to learn and generalize the distribution; the primary limiting factor for RF is architectural capacity rather than data availability.

To assess whether the generated ensembles reproduce not only the locations of dominant basins but also their configurational breadth, we compute the Shannon entropy of the marginal distributions,

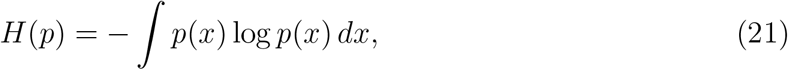

and report the entropy difference

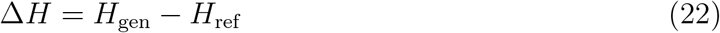

across dihedral pairs (Fig. 6). Small values of Δ*H* indicate faithful preservation of ensemble diversity, whereas positive and negative deviations correspond to over-dispersed and over-concentrated distributions, respectively.

**FIG. 6.**
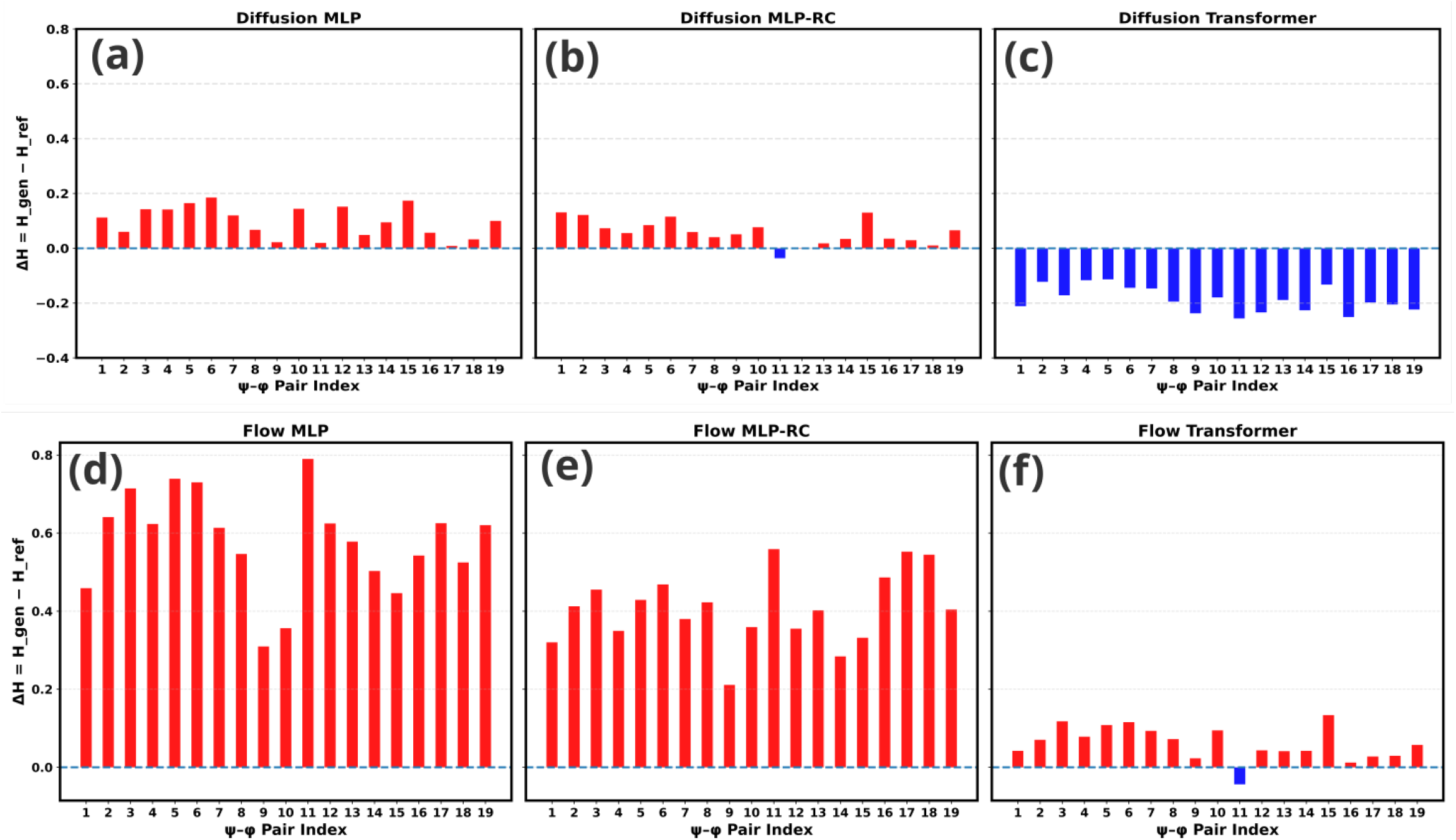
Entropy difference across Trp-cage dihedral pairs. Panels (a)–(c) correspond to diffusion models and panels (d)–(f) to rectified-flow models. Low Δ*H* indicates faithful preservation of ensemble diversity. Positive values indicate over-dispersion, whereas negative values indicate overly narrow ensembles. Diffusion, particularly with MLP-RC, preserves the reference entropy well across architectures, reflecting the stabilizing effect of stochastic relaxation on ensemble breadth. RF with MLP and MLP-RC produces strongly positive Δ*H* because insufficiently expressive velocity fields spread probability mass too broadly; only the Transformer recovers an appropriate balance between sharpness and diversity.

For diffusion, Δ*H* remains uniformly small across architectures, indicating accurate preservation of configurational diversity. The Transformer shows a slight negative Δ*H*, suggesting mild sharpening of probability mass in high-density regions. In contrast, RF with MLP and MLP-RC produces strongly positive Δ*H*, reflecting overly diffuse distributions and an overestimation of configurational entropy—a direct consequence of the velocity field being unable to confine probability mass to the correct regions when architecturally under-constrained. The Transformer substantially reduces this discrepancy, recovering a better balance between structural sharpness and ensemble diversity.

Taken together, the Trp-cage results show that diffusion remains comparatively robust to reduced architectural capacity even in a correlated protein conformational space, whereas RF becomes strongly limited by model expressivity once the target ensemble is both high-dimensional and structured.

### C. *α*-Synuclein: conformational heterogeneity amplifies the expressivity demands of RF

We finally examine whether the trends identified above persist in a broader and more heterogeneous ensemble by considering a 60-dimensional dihedral subspace of the intrinsically disordered protein *α*-synuclein, comprising 30 (*ϕ, ψ*) pairs. Unlike Trp-cage, *α*-synuclein occupies a broad conformational ensemble with weaker local confinement and substantially greater heterogeneity, making the transport problem for RF correspondingly more global and the relaxation benefit for diffusion correspondingly more pronounced.

We use a 73 *µ*s molecular dynamics trajectory from D. E. Shaw Research as the reference distribution and employ 50% of the data for training. As investigated previously by our group [26], this larger training fraction improves coverage of conformational space while retaining a nontrivial learning problem.

Representative free-energy surfaces and marginal distributions are shown in the Supporting Information (Fig. S6). Diffusion reproduces the overall structure of the conformational ensemble across architectures, with MLP-RC and Transformer providing the most consistent agreement. The plain MLP shows somewhat larger deviations but still captures the global topology of the marginals. Thus, even in this highly heterogeneous setting, diffusion remains comparatively robust to reduced architectural capacity.

In contrast, RF displays pronounced architectural dependence. The MLP and MLP-RC models generate visibly distorted marginal distributions and fail to capture the broad, correlated character of the ensemble. Only the Transformer architecture recovers the key structural features with reasonable fidelity. This behavior is quantified through KL divergence across dihedral pairs (Fig. S7): diffusion maintains relatively low KL divergence across architectures, whereas RF incurs large errors for MLP-based models, with the Transformer sharply reducing the mismatch across nearly all residues.

Importantly, this pattern persists despite the use of a larger training set, indicating that the limiting factor in RF is not merely data availability but the fundamental difficulty of representing coherent deterministic transport in a high-dimensional, heterogeneous conformational space. The *α*-synuclein results therefore strengthen and generalize the conclusion drawn from Trp-cage: as conformational heterogeneity increases, the expressivity demands placed on the deterministic transport field grow correspondingly more severe, while the stochastic relaxation mechanism of diffusion continues to provide a self-correcting buffer against architectural imperfection.

### D. Sampling dynamics reveal the mechanistic origin of the architecture gap

The endpoint analyses above establish clear differences in both sampling fidelity and architectural dependence across all three systems. We now ask whether these differences can be understood mechanistically from the sampling trajectories themselves—and show that they can, through a single unifying principle visible directly in the time evolution of the KL divergence.

A useful diagnostic is the time evolution of the KL divergence between the generated distribution *p*_*t*_(*x*) and the target distribution *π*(*x*). For diffusion processes, the evolution of KL divergence has a dissipative form,

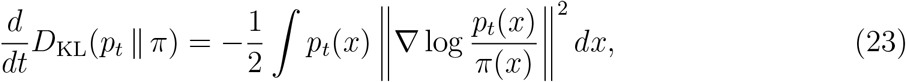

which is non-positive. This formal expression captures the fact that diffusion *intrinsically* relaxes relative entropy over time—it is not something the network needs to achieve; it is built into the dynamics. By contrast, for deterministic flow,

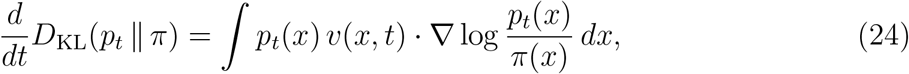

whose sign depends entirely on the learned velocity field. KL reduction is not built into deterministic transport; it must be achieved through sufficiently accurate representation of the transport field. (See SI Section III for a full convergence analysis of KL divergence over time using the Fokker–Planck equation for both paradigms.)

These theoretical differences are directly and unambiguously reflected in the sampling trajectories of the three-well potential (Fig. 7). In diffusion (Figs. 7a,b), the forward process progressively drives *p*_data_ to Gaussian noise; in the reverse process, the KL divergence remains elevated during early steps—when the distribution is still close to noise—and then drops sharply in the late stage once the denoising process resolves the basin geometry. This pronounced late-stage collapse in KL divergence is the dynamical signature of stochastic relaxation: probability mass is rapidly redistributed into the correct metastable basins as the reverse process approaches the data manifold. In RF (Fig. 7c), by contrast, the KL divergence decreases gradually and smoothly throughout, reflecting progressive deterministic transport rather than dissipative convergence. There is no late-stage drop because there is no relaxation mechanism; convergence depends entirely on the learned velocity field tracking the optimal transport path.

**FIG. 7.**
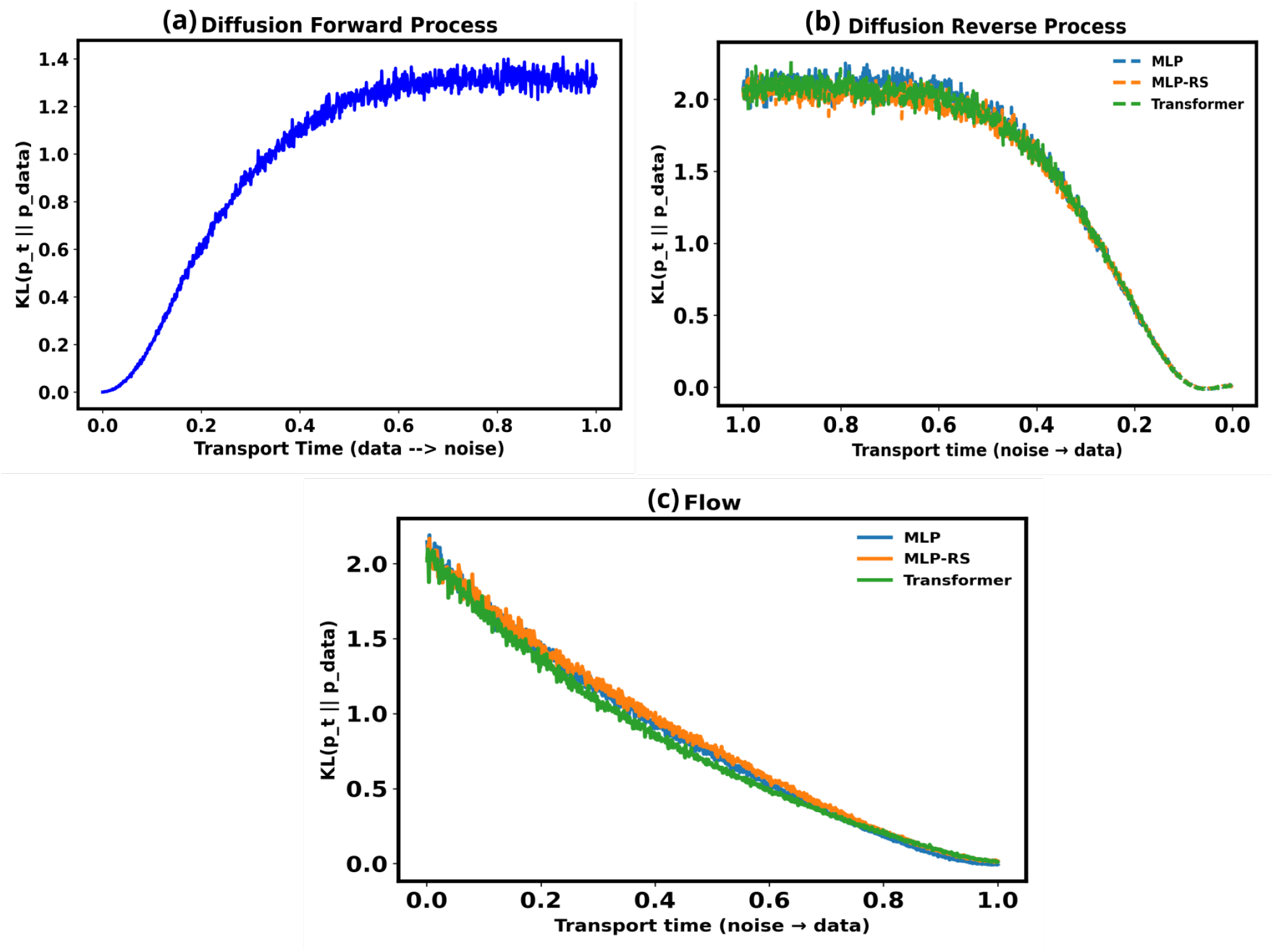
Sampling dynamics in the three-well potential reveal two mechanistically distinct convergence routes. Panel (a) shows the time evolution of KL divergence during the fixed forward diffusion process (data to noise); panel (b) shows the reverse diffusion process (noise to data). The pronounced late-stage drop in KL divergence during the reverse process is the dynamical signature of stochastic relaxation: the non-positive dissipative term in the Fokker–Planck equation guarantees monotonic contraction toward the target, and the network does not need to achieve this alone. Panel (c) shows the time evolution of KL divergence during rectified-flow sampling. RF approaches the target gradually through deterministic transport, with no late-stage collapse, because KL reduction is not built into the dynamics and must be achieved entirely through accurate velocity-field learning. The architecture dependence of RF (visible as spread between MLP, MLP-RC, and Transformer curves) is a direct consequence of this absence of a self-correcting mechanism.

To extend this mechanistic analysis to the higher-dimensional Trp-cage system, we examine the evolution of low-order moments during sampling. Direct estimation of the full probability density becomes intractable in high dimensions, so we use moment evolution as a coarse but informative proxy for convergence. For each dimension *d*, the reference mean and variance are defined as

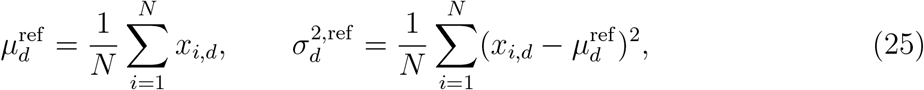

while the corresponding quantities for generated samples at sampling time *t* are

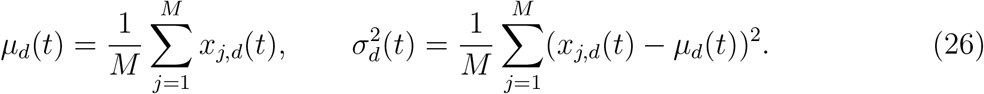

Figure 8 shows mean and variance evolution during the forward diffusion process in Trp-cage, in which the data distribution is driven to Gaussian noise: means converge to zero and variances to one, as expected from the standard Ornstein–Uhlenbeck process. This establishes the well-defined endpoint from which the reverse process begins.

**FIG. 8.**
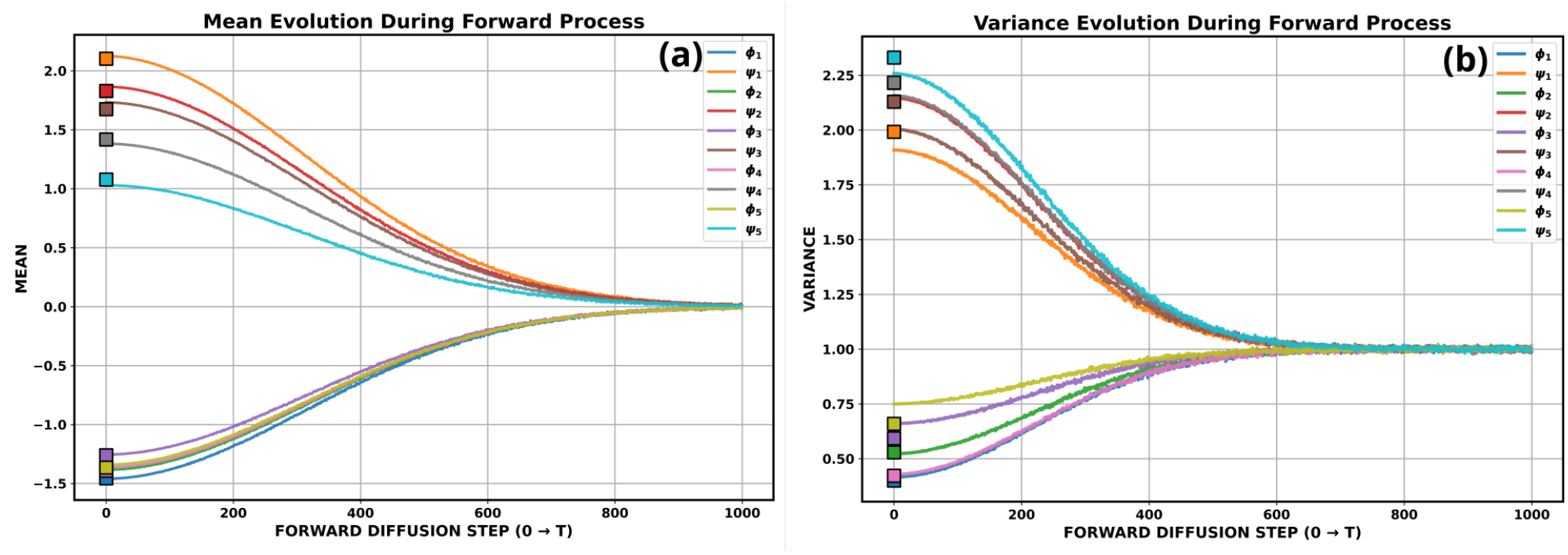
Mean and variance evolution during forward diffusion for Trp-cage. Panels (a) and (b) show the evolution of mean and variance values across selected dihedral dimensions during the forward process. Square markers indicate the reference values from the full MD ensemble. The forward process is a fixed Ornstein–Uhlenbeck dynamics that drives all means to zero and all variances to one—establishing the well-defined Gaussian endpoint from which the learned reverse process must recover the data distribution. This panel provides the dynamical baseline against which reverse-process convergence is interpreted.

Figures 9 and 10 show the evolution of these quantities during the reverse diffusion and rectified-flow processes respectively. Mean convergence reports whether probability mass is centered correctly; variance convergence reports whether configurational fluctuations are reproduced with the correct amplitude—a more sensitive probe of distributional fidelity.

**FIG. 9.**
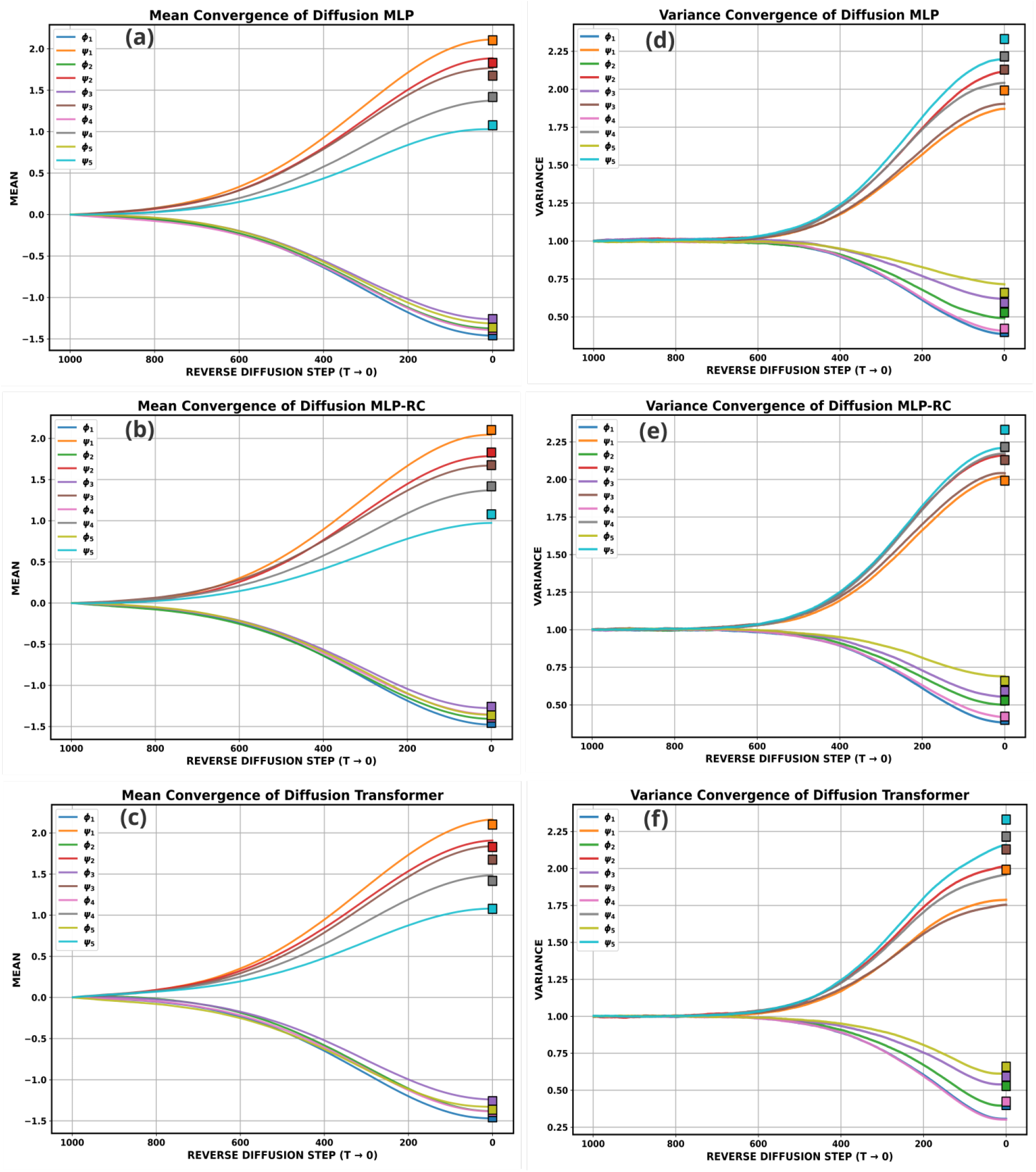
Mean and variance convergence during diffusion sampling for Trp-cage. Panels (a)–(c) show the evolution of mean values across selected dihedral dimensions, and panels (d)– (f) show the corresponding variances, for MLP, MLP-RC, and Transformer architectures. Square markers indicate the reference values from the full MD ensemble. All architectures converge to the correct means, reflecting the centering robustness conferred by stochastic relaxation. Variance convergence is most accurate for the residual architecture, indicating that residual connections pro-vide the additional stability needed to reproduce configurational fluctuation amplitudes faithfully even without Transformer-level expressivity.

**FIG. 10.**
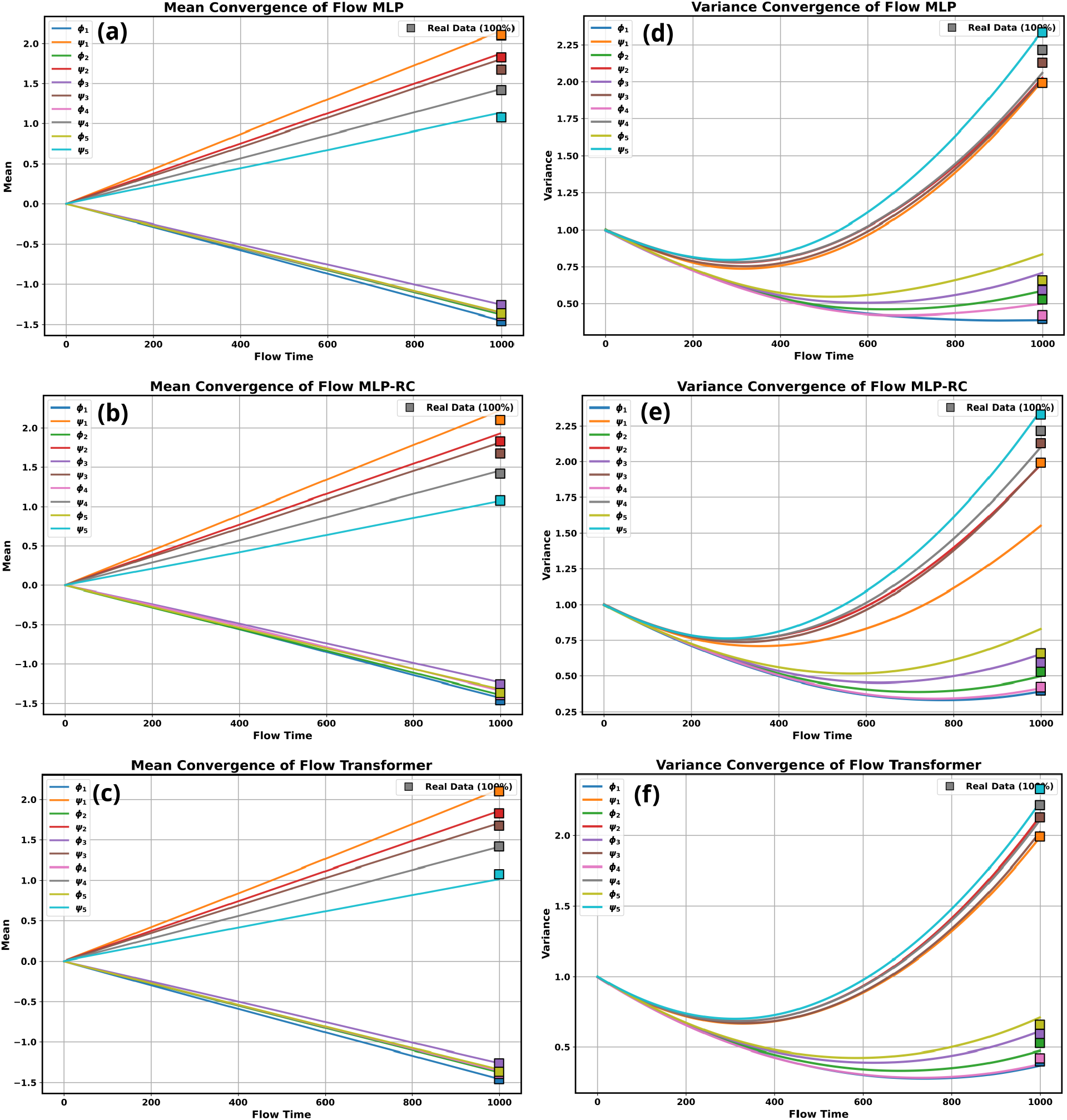
Mean and variance convergence during rectified-flow sampling for Trp-cage. Panels (a)–(c) show mean convergence and panels (d)–(f) show variance convergence for MLP, MLP-RC, and Transformer architectures. Square markers indicate the reference values from the full MD ensemble. Mean convergence follows an approximately linear trajectory in all cases—a direct dynamical signature of the linear interpolation structure of rectified flow, confirming that transport proceeds along straight-line paths without stochastic correction. Variance convergence is substantially more sensitive to architecture than in diffusion: the Transformer closely tracks the reference fluctuation amplitudes, while MLP-based models exhibit persistent deviations that are never recovered because deterministic transport provides no self-correcting mechanism for velocity-field errors.

For diffusion (Fig. 9), all architectures converge closely to the correct mean values across dimensions, and variance convergence is most accurate for the MLP-RC model. This pattern is mechanistically consistent with the KL trajectory: the stochastic reverse process centers probability mass reliably even for weaker architectures, and the residual connections provide the additional stability needed to track configurational fluctuations accurately.

For RF (Fig. 10), the architecture dependence is considerably stronger. The Transformer closely tracks both the mean and variance across dimensions, whereas MLP-based models show persistent deviations, especially in variance. Notably, mean convergence in RF follows an approximately linear trajectory—a direct consequence of the linear interpolation structure of the rectified-flow formulation, *x*_*t*_ = (1 − *t*)*z* + *tx*. This linearity is a visual confirmation that the dynamics are purely deterministic transport along straight-line paths, with no stochastic correction available. Additional dimensions are shown in the Supporting Information (Figs. S8–S12), where the same pattern holds throughout.

These dynamical diagnostics provide the mechanistic interpretation that unifies all the endpoint results. Diffusion achieves convergence through stochastic relaxation, which redistributes probability mass even when the learned model is imperfect. RF relies directly on accurate deterministic transport and therefore requires sufficient model expressivity to reproduce both the location and spread of the target distribution. When the architecture is inadequate, errors in the transport field manifest as persistent discrepancies in higher-order statistics that are never corrected—because, unlike diffusion, the dynamics provide no intrinsic recovery mechanism.

## IV. DISCUSSION

The central contribution of this work is mechanistic rather than benchmarking. Diffusion and rectified flow are not simply two methods that differ in accuracy; they embody two qualitatively distinct dynamical routes to the same target distribution, and understanding those routes changes how one should think about generative model selection, architectural design, and the role of stochasticity in molecular conformational sampling.

### Stochastic relaxation as an intrinsic convergence mechanism

The most distinctive feature of diffusion sampling is visible in the KL-divergence trajectory itself (Fig. 7): distributional error remains elevated during early reverse steps and then drops sharply in the late stage as the denoising process resolves basin geometry. This is not an artifact of parameterization—it reflects a fundamental property of stochastic dynamics. As formalized through the Fokker–Planck equation (see SI, Section III), the diffusion term introduces a strictly non-positive contribution to the time derivative of KL divergence,

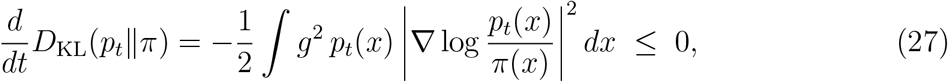

guaranteeing dissipative contraction toward equilibrium. In practice, this means the stochastic reverse process can redistribute probability mass into the correct metastable regions even when the learned score function is imperfect—the noise itself acts as a corrective regularizer. Stochasticity in diffusion is therefore not merely an implementation choice; it is a functional source of robustness that partially compensates for approximation errors in the neural network. This feature explains why diffusion remains accurate even with relatively simple architectures, and why the residual MLP often performs nearly as well as, or better than, the Transformer in the present benchmarks (Figs. 5, 6).

### Deterministic transport: efficiency contingent on expressivity

Rectified flow converges through a qualitatively different mechanism. Because the continuity equation lacks the Laplacian entropy-producing term, the time derivative of KL divergence is

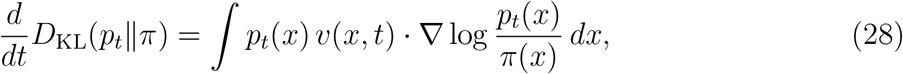

whose sign depends entirely on the learned velocity field. There is no intrinsic guarantee of monotonic KL reduction; convergence depends on how accurately *v*_*θ*_ approximates the true transport map. This is directly reflected in the KL-divergence trajectory (Fig. 7(c)): RF approaches the target gradually and smoothly, without the late-stage drop characteristic of diffusion. When the architecture is sufficiently expressive—as with the Transformer—RF recovers the target distribution with high accuracy. When it is not, errors in the transport field accumulate deterministically and manifest as persistent deviations in both mean and variance (Fig. 10), elevated KL divergence (Fig. 5), and inflated configurational entropy (Fig. 6). The consistent improvement obtained with the Transformer therefore reflects more than increased parameter count; it points to a structural necessity for global feature mixing in deterministic generative transport, a requirement that becomes more severe as the target ensemble grows broader and more correlated.

### Sampling dynamics as the primary diagnostic

A key finding of this work is that endpoint distributions alone are an insufficient characterization of generative model behavior. The convergence trajectories of diffusion and RF are mechanistically distinguishable—in their KL profiles, their moment evolution, and their sensitivity to architectural capacity—in ways that endpoint comparisons erase. Diffusion shows a characteristic late-stage reduction in KL divergence, consistent with dissipative stochastic relaxation, while RF approaches the target through transport that depends explicitly on the accuracy of the learned velocity field. The moment-evolution analysis for Trp-cage reinforces the same picture: diffusion recovers the center and spread of the distribution across architectures (Fig. 9), whereas RF exhibits persistent deviations in variance for MLP-based models (Fig. 10), deviations that are not corrected during sampling because deterministic transport provides no self-correcting mechanism. Evaluating generative models only by final sample quality would therefore miss this important mechanistic distinction. The diagnostics used here—KL divergence, Shannon entropy, and moment evolution—provide complementary windows into convergence behavior: KL divergence measures how accurately probability mass is assigned across conformational space; entropy distinguishes over-dispersed from over-concentrated ensembles; and moment evolution provides a coarse but informative view of convergence in settings where full density estimation is intractable. Together, these observables show that diffusion and RF differ not only in where they end but in how they get there.

### Architecture as a consequence of dynamics, not an independent choice

The architecture dependence observed across all three systems is not an incidental finding; it is a direct consequence of the underlying generative dynamics. Because diffusion sampling is regularized by stochastic relaxation, the neural network’s role is to provide local denoising directions within a process that already has a built-in convergence tendency. A residual MLP is often sufficient for this purpose. Rectified flow places a fundamentally greater burden on the network: it must represent a coherent global transport field, and errors anywhere in that field propagate without correction. The Transformer’s capacity for global feature mixing is therefore not merely beneficial for RF—it is structurally necessary as the conformational landscape grows more correlated and heterogeneous. This implies that architecture should not be treated as a secondary implementation choice. Instead, the generative dynamics sets the expressivity requirement, and architecture must be chosen in relation to it. Architecture, generative dynamics, and target-system complexity must be considered jointly, not independently.

### Practical implications and the cost–fidelity tradeoff

The decision framework that emerges from these findings is grounded in the mechanistic picture rather than in benchmark rankings. Figure S13 summarizes the training and sampling times for both paradigms. In the present implementation, RF is cheaper to train because it learns the transport field directly through a single forward pass per step, whereas diffusion incurs additional cost through denoising across multiple reverse steps. However, computational efficiency cannot be considered independently of sampling fidelity: RF often requires a more expressive—and therefore more expensive— architecture to achieve accuracy comparable to diffusion, so the apparent training-time advantage of RF is partially offset by the higher architectural cost it demands.

When the target ensemble is high-dimensional, strongly correlated, or only sparsely sampled—conditions common in intrinsically disordered proteins and flexible multi-domain systems—diffusion provides a robust default: its stochastic relaxation provides a margin against approximation error that makes even comparatively simple architectures viable. Rectified flow remains an attractive alternative when efficiency is paramount and a sufficiently expressive architecture is available, but its performance advantage is only realized once the expressivity condition is met. Selecting RF with an inadequate architecture not only reduces accuracy; it does so in a way that is dynamically unrecoverable, because the deterministic transport lacks a self-correcting mechanism.

### A path forward: combining stochastic robustness with deterministic efficiency

The present results should be interpreted as evidence for a broader and likely general principle: stochastic generative dynamics can partially compensate for limited expressivity, whereas deterministic transport places stricter demands on model architecture. This principle has implications beyond the specific paradigms compared here. Future progress in molecular generative modeling is unlikely to come from refining either diffusion or RF in isolation, but from developing hybrid approaches that combine their respective strengths. Methods that inject controlled stochasticity into transport-based sampling, or that learn velocity fields with built-in dissipative regularization, may offer a more favorable balance between efficiency and fidelity than either pure paradigm. Likewise, architectures explicitly designed to capture molecular correlations—through equivariance, physically informed attention patterns, or coordinate-aware representations—may reduce the expressivity burden that currently makes deterministic transport sensitive to model quality. Such developments will be essential for extending learned generative samplers to larger, more heterogeneous biomolecular systems and for bringing these methods closer to physically faithful molecular simulation.

## CODE AVAILABILITY

The details of the implementation of the model and scripts can be found at the following URL:https://github.com/nagesh123-geek/Diffusion_Flow_sampling_dynamics

## SUPPLEMENTARY INFORMATION

The supplementary information (SI) provides supplementary figures and mathematical details of the model used. We introduce the mathematical foundations of Diffusion and flow models then we derive the KL-divergence time evolution using Fokker-Planck equation to show the convergence behaviour of Diffusion and Flow models(Section I to Section V).Then we show the minimal ELBO for DDPM and Flow models(Section VI). In section VII we show the mathematical details of architectures used for training. In section VIII, we show the relationship between KL divergence and entropy which we used in analysis of data sampled. As additional proof of concept we show the FES of different *ψ* − *ϕ* angles of trpcage in Fig S1, S2 and S3 for both diffusion and flow models. In figure S4 we show the Error study of KL divergence across selected Trpcage dihedral pairs using 10% training data using 10 sampled samples. In Fig S5, KL divergence is computed across selected Trpcage dihedral pairs using 30% data as training data. This increasing data is to account for the minimal data needed capture the distribution for modelling and to see what happens if we increase the training data. In Fig S6, Representative (*ψ*), (*ϕ*) marginals for *α*−synuclein are shown. In Fig S7 corresponding KL divergence across selected *α*−synuclein dihedral pairs are shown. In Fig S8, we show the Mean and variance convergence to noise during forward diffusion process applied on *ϕ*_6_, *ψ*_6_, *ϕ*_7_, *ψ*_7_, *ϕ*_8_, *ψ*_8_, *ϕ*_9_, *ψ*_9_, *ϕ*_10_, *ψ*_10_, *ϕ*_11_, *ψ*_11_, *ϕ*_12_, *ψ*_12_, *ϕ*_13_, *ψ*_13_, *ϕ*_14_, *ψ*_14_, *ϕ*_15_, *ψ*_15_ for Trp-cage.

Fig S9 shows mean and variance convergence during diffusion sampling for Trp-cage of *ϕ*_6_, *ψ*_6_, *ϕ*_7_, *ψ*_7_, *ϕ*_8_, *ψ*_8_, *ϕ*_9_, *ψ*_9_, *ϕ*_10_, *ψ*_10_.

Fig S10 shows mean and variance convergence during rectified-flow sampling for Trp-cage of *ϕ*_6_, *ψ*_6_, *ϕ*_7_, *ψ*_7_, *ϕ*_8_, *ψ*_8_, *ϕ*_9_, *ψ*_9_, *ϕ*_10_, *ψ*_10_.

Fig S11 shows the mean and variance convergence during diffusion sampling for Trp-cage of *ϕ*_11_, *ψ*_11_, *ϕ*_12_, *ψ*_12_, *ϕ*_13_, *ψ*_13_, *ϕ*_14_, *ψ*_14_, *ϕ*_15_, *ψ*_15_.

Fig S12 shows the mean and variance convergence during rectified-flow sampling for Trp-cage of *ϕ*_11_, *ψ*_11_, *ϕ*_12_, *ψ*_12_, *ϕ*_13_, *ψ*_13_, *ϕ*_14_, *ψ*_14_, *ϕ*_15_, *ψ*_15_.

Fig S13 shows the training and sampling time for diffusion and rectified-flow models.

## ACKNOWLEDGEMENT

We acknowledge support of the Department of Atomic Energy, Government of India, under Project Identification No. RTI 4007. All the authors acknowledge Tata Institute of Fundamental Research Hyderabad, India for providing the support of computing resources. JM acknowledges funding from Department of Science and Technology of India (CRG/2023/001426).

