## Supplemental Information for "How Generative Models Approach Molecular Conformational Sampling"

Nagesh B E[*] and Jagannath Mondal[†]

*Tata Institute of Fundamental Research, 36/P,*

*Gopanpally Village, Serilingampally Mandal,*

*Ranga Reddy District, Hyderabad 500046, Telangana, India*

(Dated: April 10, 2026)

### I. MATHEMATICAL THEORY

#### Theory of Diffusion and Flow Models

Diffusion models are stochastic differential equation (SDE)–based generative models that explicitly learn probability distributions through two stochastic processes: a forward process and a reverse process. In the forward process, no learning is involved; instead, noise is progressively added to the data according to a predefined variance schedule $\beta$ over a fixed time horizon T . This process gradually transforms the data distribution into a simple reference distribution (typically Gaussian noise) and, in the continuous-time limit, can be viewed as an Ornstein–Uhlenbeck (OU)–type process. The reverse process, in contrast, is learned. A neural network parameterization is trained to approximate the time-reversed dynamics, effectively learning to denoise samples and reconstruct the data distribution starting from pure noise. Through this learned reverse diffusion, the model generates samples from the target distribution. Flow models, on the other hand, are ordinary differential equation (ODE)–based generative models that explicitly learn probability distributions via deterministic transport. Rather than relying on stochastic perturbations, flow models learn a time-dependent velocity field that continuously transports samples from a base distribution to the target data distribution. This transport can be interpreted through the lens of optimal transport theory, where the learned dynamics define a mapping between the two distributions through deterministic trajectories in state space.

[*]

[†]

### II. UNIFIED DERIVATION OF DIFFUSION AND FLOW MODELS FROM THE FOKKER–PLANCK EQUATION

#### A. General Density Evolution

The time evolution of a probability density $p_t(x)$ associated with a stochastic process

$$dx_t = u(x_t, t)\,dt + \sqrt{D(x_t, t)}\,dW_t \tag{1}$$

is governed by the Fokker–Planck equation:

$$\partial_t p_t(x) = -\nabla \cdot \big(p_t(x)u(x, t)\big) + \frac{1}{2}\nabla^2\big(D(x, t)p_t(x)\big), \tag{2}$$

where $u(x, t)$ is the drift term and $D(x, t)$ is the diffusion coefficient.

Equation (2) describes the most general continuous-time density evolution combining deterministic transport and stochastic diffusion.

#### B. Diffusion Models (DDPM) from Fokker–Planck

Diffusion models define a forward stochastic differential equation (SDE):

$$dx_t = f(x_t, t)\,dt + g(t)\,dW_t. \tag{3}$$

For the variance-preserving formulation,

$$f(x, t) = -\frac{1}{2}\beta(t)x, \qquad g(t)^2 = \beta(t), \tag{4}$$

which corresponds to a time-dependent Ornstein–Uhlenbeck process.

Substituting into the Fokker–Planck equation gives:

$$\partial_t p_t = -\nabla \cdot \left(-\frac{1}{2}\beta(t)x\,p_t\right) + \frac{1}{2}\beta(t)\nabla^2 p_t. \tag{5}$$

Simplifying,

$$\partial_t p_t = \frac{1}{2}\beta(t)\nabla \cdot (xp_t) + \frac{1}{2}\beta(t)\nabla^2 p_t. \tag{6}$$

As $t \to T$, the density converges to a Gaussian distribution,

$$p_T(x) \to \mathcal{N}(0, I). \tag{7}$$

*Reverse-Time SDE*

The reverse-time dynamics (Anderson, 1982) are given by:

$$dx_t = \left[f(x,t) - g(t)^2 \nabla \log p_t(x)\right] dt + g(t) d\bar{W}_t. \tag{8}$$

For the variance-preserving case,

$$dx_t = \left[-\frac{1}{2}\beta(t)x - \beta(t)\nabla \log p_t(x)\right] dt + \sqrt{\beta(t)}\, d\bar{W}_t. \tag{9}$$

Diffusion models train a neural network to approximate the score function:

$$\nabla \log p_t(x). \tag{10}$$

Thus, DDPM arises directly from the Fokker–Planck equation combined with reverse-time SDE theory.

#### C. Flow Models from the Continuity Equation

If we remove stochastic diffusion by setting

$$D(x,t) = 0, \tag{11}$$

Equation (2) reduces to

$$\partial_t p_t(x) = -\nabla \cdot \big(p_t(x) v(x,t)\big), \tag{12}$$

which is the continuity equation.

The corresponding dynamics are governed by the deterministic ODE:

$$\frac{dx}{dt} = v(x,t). \tag{13}$$

Flow-based generative models learn the velocity field $v(x,t)$ that transports samples from a base distribution $p_0$ to the target distribution $p_T$.

#### D. Normalizing Flows via Change of Variables

Normalizing flows (NF) model the transformation from a simple base distribution $p_0(z)$ to a complex target distribution $p(x)$ using a sequence of invertible mappings. Let $x = f(z)$, where $f$ is a bijective transformation. Then, the density of $x$ is given by the change-of-variables formula:

$$p(x) = p_0(z) \left| \det \left( \frac{\partial f^{-1}(x)}{\partial x} \right) \right|. \tag{14}$$

For a composition of $K$ invertible transformations,

$$x = f_K \circ f_{K-1} \circ \cdots \circ f_1(z), \tag{15}$$

the log-density can be written as

$$\log p(x) = \log p_0(z) - \sum_{k=1}^{K} \log \left| \det \left( \frac{\partial f_k(x_{k-1})}{\partial x_{k-1}} \right) \right|, \tag{16}$$

where $x_0 = z$ and $x_K = x$.

The requirement of exact likelihood evaluation necessitates that each transformation be invertible and have a tractable Jacobian determinant. This constraint restricts the choice of architectures and becomes increasingly expensive in high-dimensional settings due to the computation of determinants.

In contrast to flow models derived from the continuity equation, which learn a velocity field $v(x, t)$ and evolve samples through a continuous-time ODE, normalizing flows rely on discrete, invertible mappings with exact density tracking at each step. While NF provides exact likelihoods, this comes at the cost of architectural constraints and reduced flexibility in modeling complex high-dimensional distributions.

#### E. Probability Flow ODE: The Bridge Between Diffusion and Flow

For the stochastic SDE in Equation (3), there exists a deterministic ODE that produces identical marginal densities. This is known as the probability flow ODE:

$$\frac{dx}{dt} = f(x, t) - \frac{1}{2} g(t)^2 \nabla \log p_t(x). \tag{17}$$

Substituting this velocity field into the continuity equation (12) yields the same density evolution as the stochastic Fokker–Planck equation.

Thus, diffusion and flow models share the same marginal density path, but differ in their dynamical realizations:

- Diffusion models use stochastic dynamics (SDE).
- Flow models use deterministic transport (ODE).

#### F. Unified Perspective

The general density evolution equation

$$\partial_t p_t = -\nabla \cdot (p_t v) + \frac{1}{2}\nabla^2 (D p_t) \tag{18}$$

provides a unified framework:

- If $D(x,t) = 0$, the model reduces to deterministic flow.
- If $D(x,t) > 0$, the model corresponds to diffusion.

The additional Laplacian term in diffusion,

$$\frac{1}{2}\nabla^2 (D p_t), \tag{19}$$

introduces entropy production and smoothing of the density, which can enhance convergence properties relative to purely deterministic flows.

Therefore, diffusion and flow models can be interpreted as two distinct dynamical realizations of probability transport governed by the Fokker–Planck framework.

### III. CONVERGENCE OF THE PROBABILITY DISTRIBUTION OVER TIME

#### A. Preliminaries

Let $p_t(x)$ denote the time-evolving model density and let $\pi(x)$ denote the target distribution. We analyze convergence through the Kullback–Leibler (KL) divergence:

$$\mathrm{KL}(p_t \| \pi) = \int p_t(x) \log \frac{p_t(x)}{\pi(x)}\, dx. \tag{20}$$

The time derivative of the KL divergence is

$$\frac{d}{dt}\mathrm{KL}(p_t \| \pi) = \int \partial_t p_t(x) \left( \log \frac{p_t(x)}{\pi(x)} + 1 \right) dx. \tag{21}$$

We now analyze this quantity for diffusion and flow dynamics separately.

### B. Convergence Under Diffusion Dynamics (SDE)

Consider the stochastic differential equation

$$dx_t = f(x_t, t)\,dt + g(t)\,dW_t, \tag{22}$$

whose density evolves according to the Fokker–Planck equation:

$$\partial_t p_t = -\nabla \cdot (f p_t) + \frac{1}{2}\nabla^2(g^2 p_t). \tag{23}$$

Substituting Equation (23) into Equation (21):

$$\frac{d}{dt}\mathrm{KL} = \int \left[-\nabla \cdot (f p_t) + \frac{1}{2}\nabla^2(g^2 p_t)\right]\left(\log \frac{p_t}{\pi} + 1\right) dx. \tag{24}$$

*Drift Term*

Using integration by parts:

$$\int -\nabla \cdot (f p_t)\,\phi\,dx = \int f p_t \cdot \nabla\phi\,dx, \tag{25}$$

where $\phi = \log \frac{p_t}{\pi} + 1$.

*Diffusion Term*

Again using integration by parts:

$$\int \frac{1}{2}\nabla^2(g^2 p_t)\,\phi\,dx = -\frac{1}{2}\int g^2 p_t\,|\nabla\phi|^2\,dx. \tag{26}$$

Putting everything together:

$$\frac{d}{dt}\mathrm{KL}(p_t\|\pi) = \int f p_t \cdot \nabla \log \frac{p_t}{\pi}\,dx - \frac{1}{2}\int g^2 p_t \left|\nabla \log \frac{p_t}{\pi}\right|^2 dx. \tag{27}$$

If the drift $f$ is chosen so that $\pi$ is stationary, then the first term cancels and we obtain

$$\boxed{\frac{d}{dt}\mathrm{KL}(p_t\|\pi) = -\frac{1}{2}\int g^2 p_t \left|\nabla \log \frac{p_t}{\pi}\right|^2 dx} \tag{28}$$

This shows:

- KL divergence decreases monotonically.
- Convergence is strictly dissipative.
- The decay rate is controlled by the Fisher information.

Thus diffusion induces entropy production and guarantees contraction toward equilibrium.

#### C. Convergence Under Flow Dynamics (ODE)

Now consider deterministic flow:

$$\frac{dx}{dt} = v(x, t), \tag{29}$$

with density evolution governed by the continuity equation:

$$\partial_t p_t = -\nabla \cdot (p_t v). \tag{30}$$

Substituting into the KL derivative:

$$\frac{d}{dt}\mathrm{KL} = \int -\nabla \cdot (p_t v) \left(\log \frac{p_t}{\pi} + 1\right) dx. \tag{31}$$

Using integration by parts:

$$\boxed{\frac{d}{dt}\mathrm{KL}(p_t \| \pi) = \int p_t v \cdot \nabla \log \frac{p_t}{\pi}\, dx.} \tag{32}$$

Unlike diffusion, there is no negative quadratic term. The sign of the KL derivative depends entirely on the learned velocity field $v$.

Therefore:

- Convergence is not automatically dissipative.
- KL may decrease, increase, or oscillate.
- Stability depends on how well $v$ approximates optimal transport.

#### D. Comparison of Convergence Mechanisms

- Diffusion contains a Laplacian term producing

$$-\frac{1}{2}\int g^2 p_t \left|\nabla \log \frac{p_t}{\pi}\right|^2 dx,$$

  which ensures monotonic KL decay.

- Flow models lack this entropy-producing term and rely purely on deterministic transport.

- The stochastic smoothing in diffusion promotes stronger contraction toward equilibrium.

#### E. Conclusion

Diffusion models exhibit guaranteed dissipative convergence under suitable conditions due to the presence of the diffusion (Laplacian) term in the Fokker–Planck equation. In contrast, flow models evolve densities through deterministic transport governed by the continuity equation, and their convergence behavior depends entirely on the learned velocity field.

Thus, diffusion and flow models represent two fundamentally different convergence mechanisms within the unified Fokker–Planck framework.

### IV. EXPONENTIAL CONVERGENCE OF LANGEVIN PROCESSES

#### A. Overdamped Langevin Dynamics

Let the target distribution be

$$\pi(x) = \frac{1}{Z} e^{-V(x)}, \tag{33}$$

where $V : \mathbb{R}^d \to \mathbb{R}$ is a smooth potential and $Z$ is the normalizing constant.

The overdamped Langevin SDE is:

$$dx_t = -\nabla V(x_t)\, dt + \sqrt{2}\, dW_t. \tag{34}$$

Its density $p_t(x)$ evolves according to the Fokker–Planck equation:

$$\partial_t p_t = \nabla \cdot (p_t \nabla V) + \nabla^2 p_t. \tag{35}$$

The stationary distribution of this PDE is $\pi(x)$.

#### B. Time Derivative of KL Divergence

Define the KL divergence:

$$\mathrm{KL}(p_t \| \pi) = \int p_t \log \frac{p_t}{\pi} dx. \tag{36}$$

Taking the time derivative:

$$\frac{d}{dt}\mathrm{KL} = \int \partial_t p_t \left( \log \frac{p_t}{\pi} + 1 \right) dx. \tag{37}$$

Substitute the Fokker–Planck equation:

$$\frac{d}{dt}\mathrm{KL} = \int \left[ \nabla \cdot (p_t \nabla V) + \nabla^2 p_t \right] \left( \log \frac{p_t}{\pi} + 1 \right) dx. \tag{38}$$

Using integration by parts twice, we obtain:

$$\boxed{\frac{d}{dt}\mathrm{KL}(p_t \| \pi) = - \int p_t \left| \nabla \log \frac{p_t}{\pi} \right|^2 dx.} \tag{39}$$

Define the relative Fisher information:

$$\mathcal{I}(p_t \| \pi) = \int p_t \left| \nabla \log \frac{p_t}{\pi} \right|^2 dx. \tag{40}$$

Thus,

$$\frac{d}{dt}\mathrm{KL} = -\mathcal{I}(p_t \| \pi). \tag{41}$$

This shows monotonic decay of KL divergence.

#### C. Log-Sobolev Inequality

Assume $V(x)$ is strongly convex:

$$\nabla^2 V(x) \succeq mI \quad \text{for some } m > 0. \tag{42}$$

Then $\pi$ satisfies a log-Sobolev inequality:

$$\mathrm{KL}(p\|\pi) \leq \frac{1}{2m}\mathcal{I}(p\|\pi). \tag{43}$$

Equivalently,

$$\mathcal{I}(p\|\pi) \geq 2m\,\mathrm{KL}(p\|\pi). \tag{44}$$

#### D. Exponential Decay of KL Divergence

Combining:

$$\frac{d}{dt}\mathrm{KL} = -\mathcal{I}(p_t\|\pi) \leq -2m\,\mathrm{KL}(p_t\|\pi). \tag{45}$$

This gives the differential inequality:

$$\frac{d}{dt}\mathrm{KL} \leq -2m\,\mathrm{KL}. \tag{46}$$

Applying Grönwall's inequality:

$$\boxed{\mathrm{KL}(p_t\|\pi) \leq e^{-2mt}\mathrm{KL}(p_0\|\pi).} \tag{47}$$

Thus the Langevin process converges exponentially fast to equilibrium in KL divergence.

#### E. Convergence in Wasserstein Distance

Using Talagrand's inequality:

$$W_2^2(p_t, \pi) \leq \frac{2}{m}\mathrm{KL}(p_t\|\pi), \tag{48}$$

we obtain exponential convergence in Wasserstein distance:

$$W_2(p_t, \pi) \le Ce^{-mt}. \tag{49}$$

#### F. Interpretation

- The diffusion term generates entropy production.
- Strong convexity of $V$ ensures log-Sobolev inequality.
- KL divergence decays exponentially.
- Convergence rate is governed by curvature $m$.

This explains why Langevin-type diffusion processes exhibit stable and rapidly contracting convergence behavior, in contrast to deterministic flow models that lack entropy production.

### V. CONVERGENCE OF RECTIFIED FLOW WITH LINEAR INTERPOLATION

#### A. Rectified Flow Construction

Rectified Flow defines a deterministic transport between a base distribution $p_0(x)$ and a target distribution $p_1(x)$ using linear interpolation in data space:

$$x_t = (1-t)x_0 + tx_1, \qquad t \in [0,1], \tag{50}$$

where

$$x_0 \sim p_0, \qquad x_1 \sim p_1.$$

The velocity field is defined as

$$v(x_t, t) = x_1 - x_0. \tag{51}$$

The goal of rectified flow is to learn the conditional expectation:

$$v_\theta(x,t) \approx \mathbb{E}[x_1 - x_0 \mid x_t = x]. \tag{52}$$

The density $p_t(x)$ evolves according to the continuity equation:

$$\partial_t p_t(x) = -\nabla \cdot (p_t(x) v(x,t)). \tag{53}$$

Since there is no diffusion term, the dynamics are purely deterministic.

#### B. Time Derivative of KL Divergence

Let $\pi(x) = p_1(x)$ denote the target distribution. The KL divergence is

$$\mathrm{KL}(p_t \| \pi) = \int p_t(x) \log \frac{p_t(x)}{\pi(x)} dx. \tag{54}$$

Taking the time derivative:

$$\frac{d}{dt}\mathrm{KL} = \int \partial_t p_t \left( \log \frac{p_t}{\pi} + 1 \right) dx. \tag{55}$$

Substitute the continuity equation:

$$\frac{d}{dt}\mathrm{KL} = \int -\nabla \cdot (p_t v) \left( \log \frac{p_t}{\pi} + 1 \right) dx. \tag{56}$$

Using integration by parts:

$$\boxed{\frac{d}{dt}\mathrm{KL}(p_t \| \pi) = \int p_t(x)\, v(x,t) \cdot \nabla \log \frac{p_t(x)}{\pi(x)}\, dx.} \tag{57}$$

#### C. Behavior Under Linear Interpolation

For linear interpolation:

$$x_t = (1-t)x_0 + t x_1, \tag{58}$$

the velocity is constant along each trajectory:

$$\frac{dx_t}{dt} = x_1 - x_0. \tag{59}$$

Thus:

- Trajectories are straight lines.
- No stochastic smoothing is present.

- No entropy is produced during evolution.

Unlike diffusion models, there is no quadratic negative term such as

$$-\int p_t|\nabla \log \frac{p_t}{\pi}|^2 dx.$$

Therefore:

- KL decay is not guaranteed to be monotonic.
- Convergence depends entirely on how well $v_\theta$ approximates the true transport field.
- Errors in the velocity field accumulate deterministically.

#### D. Exact Convergence Under Perfect Velocity

If the learned velocity equals the optimal transport velocity:

$$v(x,t) = \mathbb{E}[x_1 - x_0 \mid x_t = x], \tag{60}$$

then the induced flow map $T_t$ satisfies:

$$p_t = (T_t)_\# p_0, \tag{61}$$

and at $t = 1$:

$$p_1 = \pi. \tag{62}$$

Thus:

$$\mathrm{KL}(p_1 \| \pi) = 0. \tag{63}$$

Convergence is exact but not necessarily dissipative during intermediate times.

#### E. Comparison with Diffusion

For diffusion models:

$$\frac{d}{dt}\mathrm{KL} = -\frac{1}{2}\int g^2 p_t \left|\nabla \log \frac{p_t}{\pi}\right|^2 dx \leq 0. \tag{64}$$

This guarantees monotonic KL contraction.

For rectified flow:

$$\frac{d}{dt}\mathrm{KL} = \int p_t v \cdot \nabla \log \frac{p_t}{\pi} dx, \tag{65}$$

which does not have a definite sign.

#### F. Conclusion

Rectified flow with linear interpolation transports probability mass deterministically along straight trajectories. Convergence to the target distribution is exact if the velocity field is learned perfectly, but unlike diffusion models, the dynamics do not contain an intrinsic entropy-producing term. As a result, convergence is gradual and transport-driven rather than dissipative, which explains the smoother and more progressive convergence behavior often observed in practice.

### VI. ELBO FOR DENOISING DIFFUSION AND RECTIFIED FLOWS

##### *1. DDPM*

Let the forward diffusion process be

$$q(x_{1:T} \mid x_0) = \prod_{t=1}^{T} q(x_t \mid x_{t-1}), \tag{66}$$

and the reverse generative process be

$$p_\theta(x_{0:T}) = p(x_T) \prod_{t=1}^{T} p_\theta(x_{t-1} \mid x_t), \tag{67}$$

where $p(x_T)$ is a simple prior (e.g., Gaussian).

The log-likelihood of the data can be lower bounded as:

$$\log p_\theta(x_0) \geq \mathbb{E}_{q(x_{1:T}|x_0)}\left[\log \frac{p_\theta(x_{0:T})}{q(x_{1:T} \mid x_0)}\right] = \mathcal{L}_{\text{ELBO}}. \tag{68}$$

Expanding the ELBO:

$$\mathcal{L}_{\text{ELBO}} = \mathbb{E}_{q(x_{1:T}|x_0)}\left[\log p_\theta(x_T) + \sum_{t=1}^{T} \log p_\theta(x_{t-1} \mid x_t) - \log q(x_t \mid x_{t-1})\right]. \tag{69}$$

For Gaussian forward and reverse processes, this reduces to a sum of mean-squared-error terms:

$$\mathcal{L}_{\text{ELBO}} \approx \sum_{t=1}^{T} \mathbb{E}_{x_0,\epsilon\sim\mathcal{N}(0,I)}\left[\frac{\beta_t^2}{2\sigma_t^2}\|\epsilon - \epsilon_\theta(x_t,t)\|^2\right], \tag{70}$$

where $\epsilon_\theta(x_t, t)$ predicts the noise added at step $t$.

*2. Rectified Flows (RF)*

For deterministic flows defined by a velocity field $v_\theta(x, t)$:

$$\frac{dx}{dt} = v_\theta(x,t), \quad x(0) \sim p_0, \tag{71}$$

the likelihood of the target $x_T$ can be expressed via the change of variables formula:

$$\log p_\theta(x_T) = \log p_0(x_0) - \int_0^T \nabla \cdot v_\theta(x(t),t)\, dt, \tag{72}$$

where the divergence term accounts for the change in volume along the flow.

This can be interpreted as an ELBO for deterministic transport:

$$\mathcal{L}_{\text{RF}} = \mathbb{E}_{x_0\sim p_0}\left[\log p_0(x_0) - \int_0^T \nabla \cdot v_\theta(x(t),t)\, dt\right]. \tag{73}$$

In contrast to DDPM, the RF ELBO does not involve stochastic noise, and the learning objective directly maximizes the transported density from the base to the target distribution.

### VII. NEURAL ARCHITECTURES FOR TIME-DEPENDENT VECTOR FIELD LEARNING

All three architectures parameterize a time-dependent function

$$f_\theta : \mathbb{R}^d \times [0,1] \to \mathbb{R}^d, \tag{74}$$

where $d = 38$ in our case. Depending on the model, $f_\theta$ represents:

- $\epsilon_\theta(x,t)$ for DDPM (noise prediction),
- $s_\theta(x,t)$ for score-based diffusion,
- $v_\theta(x,t)$ for flow / rectified flow models.

All architectures incorporate a learnable time embedding and output a vector of dimension $d$.

#### A. 1. Multi-Layer Perceptron (MLP)

##### 1. *Time Embedding*

Time $t \in \mathbb{R}$ is mapped to a higher-dimensional representation:

$$\phi(t) = \sigma\left(W_2\sigma(W_1 t + b_1) + b_2\right), \quad \phi(t) \in \mathbb{R}^h, \tag{75}$$

where $h = 256$ and $\sigma$ denotes ReLU activation.

##### 2. *Concatenation*

The input is augmented by concatenating data and time embedding:

$$h_0 = \begin{bmatrix} x \\ \phi(t) \end{bmatrix} \in \mathbb{R}^{d+h}. \tag{76}$$

##### 3. *Hidden Layers*

The MLP consists of four blocks of the form:

$$h_{k+1} = \sigma\big(\mathrm{LN}(W_k h_k + b_k)\big), \tag{77}$$

where LN denotes Layer Normalization.

*4. Output Layer*

$$f_\theta(x,t) = W_{\text{out}} h_4 + b_{\text{out}}. \tag{78}$$

*5. Interpretation*

The MLP implements a nonlinear map:

$$f_\theta(x,t) = \text{MLP}_\theta\big([x,\phi(t)]\big), \tag{79}$$

treating time as an additional feature dimension.

### B. 2. Residual MLP (MLP-RC)

The residual variant introduces skip connections.

*1. Initial Projection*

$$h_0 = \sigma\left(\text{LN}(W_{\text{in}}[x,\phi(t)] + b_{\text{in}})\right). \tag{80}$$

*2. Residual Blocks*

Each residual block computes:

$$F(h) = \sigma\left(\text{LN}\big(W_2\sigma(\text{LN}(W_1 h))\big)\right), \tag{81}$$

and updates:

$$h_{k+1} = h_k + F(h_k). \tag{82}$$

After four residual blocks:

$$h_4 = h_0 + \sum_{k=1}^{4} F_k(h_{k-1}). \tag{83}$$

*3. Output*

$$f_\theta(x,t) = W_{\text{out}} h_4 + b_{\text{out}}. \tag{84}$$

*4. Interpretation*

Residual connections approximate a discretized dynamical system:

$$\frac{dh}{ds} = F(h), \tag{85}$$

making the network structurally aligned with continuous-time dynamics. This improves gradient flow and stability for vector field learning.

#### C. 3. Transformer-Based Model

*1. Input Projection*

$$h_0 = \text{LN}(W_{\text{proj}} x). \tag{86}$$

*2. Time Conditioning (Additive)*

Instead of concatenation, time is added:

$$h_1 = h_0 + \phi(t). \tag{87}$$

This corresponds to FiLM-style additive conditioning.

*3. Self-Attention Encoder*

The hidden state is reshaped as a sequence of length one:

$$h_1 \in \mathbb{R}^{1\times h}. \tag{88}$$

Each transformer layer performs:

$$\tilde{h} = h + \text{SelfAttention}(h), \tag{89}$$

$$h' = \tilde{h} + \text{FeedForward}(\tilde{h}), \tag{90}$$

with Layer Normalization applied in pre-norm configuration.

Since the sequence length is one, self-attention simplifies to learned linear mixing within the feature space.

##### *4. Output Projection*

$$f_\theta(x, t) = W_2 \sigma(W_1 h_L), \tag{91}$$

where $L$ is the number of encoder layers.

##### *5. Self-Attention Mechanism*

For a hidden representation $h \in \mathbb{R}^{n \times d}$, self-attention operates by first projecting $h$ into query, key, and value spaces:

$$Q = hW_Q, \qquad K = hW_K, \qquad V = hW_V, \tag{92}$$

where $W_Q, W_K, W_V \in \mathbb{R}^{d \times d_k}$ are learnable projection matrices.

The attention weights are computed as

$$A = \text{softmax}\left(\frac{QK^\top}{\sqrt{d_k}}\right), \tag{93}$$

and the output of the attention layer is

$$\text{SelfAttention}(h) = AV. \tag{94}$$

In the multi-head setting with $H$ heads, the outputs from different heads are concatenated and linearly projected:

$$\text{MHA}(h) = \text{Concat}(\text{head}_1, \ldots, \text{head}_H) W_O. \tag{95}$$

*a. Special Case: Sequence Length One* In our setting, the input is reshaped as $h \in \mathbb{R}^{1\times d}$. Consequently,

$$QK^\top \in \mathbb{R}^{1\times 1}, \tag{96}$$

and the softmax reduces to

$$A = 1. \tag{97}$$

Thus, self-attention simplifies to

$$\text{SelfAttention}(h) = V = hW_V, \tag{98}$$

which corresponds to a learned linear transformation within the feature space. In the multi-head case, this results in a richer but still purely feature-wise mixing without any token-to-token interaction.

#### 6. *Interpretation*

The Transformer introduces:

- Multi-head feature mixing,
- Deeper residual pathways,
- Stronger representational capacity.

Even with sequence length one, the stacked attention blocks act as structured residual transformations.

### D. Number of Trainable Parameters

Even though all three models use the same hidden dimension, the number of trainable parameters differs significantly due to differences in architectural complexity.

*a. MLP (Multi-Layer Perceptron)* The MLP is the simplest architecture, consisting only of fully connected layers. The number of parameters mainly comes from the weight matrices between layers. Since there are no additional components such as attention or conditioning mechanisms, the total parameter count remains relatively low.

*b. MLP-RC (MLP with Residual Connections)* The MLP-RC model introduces additional conditioning mechanisms. This requires extra layers or projections to process the conditioning inputs. As a result, more weight matrices are added to the model, increasing the total number of trainable parameters compared to the standard MLP, even though the hidden dimension remains the same.

*c. Transformer* The Transformer architecture has the highest number of parameters due to its use of multi-head self-attention and deeper structure. Each Transformer layer includes multiple projection matrices for query, key, and value computations, as well as feed-forward networks. Additionally, stacking multiple layers further increases the parameter count. Thus, even with the same hidden dimension, the use of attention mechanisms and layer depth leads to a significantly larger number of parameters.

| **Model Type** | **Number of Parameters** |
|---|---|
| MLP | 218406 |
| MLP-RC | 682534 |
| Transformer | 4890406 |

TABLE I. Model types and their number of parameters

### VIII. RELATIONSHIP BETWEEN ENTROPY AND KULLBACK–LEIBLER DIVERGENCE

Let $P(x)$ and $Q(x)$ be two probability distributions defined over the same support.

#### A. Shannon Entropy

The Shannon entropy of a distribution $P$ is defined as

$$H(P) = -\int P(x) \log P(x) \, dx. \tag{99}$$

Entropy measures the uncertainty or randomness of a distribution.

#### B. Cross-Entropy

The cross-entropy between $P$ and $Q$ is defined as

$$H(P, Q) = -\int P(x) \log Q(x)\, dx. \tag{100}$$

#### C. Kullback–Leibler Divergence

The Kullback–Leibler (KL) divergence from $Q$ to $P$ is defined as

$$D_{\mathrm{KL}}(P\|Q) = \int P(x) \log \frac{P(x)}{Q(x)}\, dx. \tag{101}$$

#### D. Relationship

Expanding the logarithm, we obtain

$$D_{\mathrm{KL}}(P\|Q) = \int P(x) \log P(x)\, dx - \int P(x) \log Q(x)\, dx \tag{102}$$

$$= -H(P) + H(P, Q). \tag{103}$$

Therefore,

$$D_{\mathrm{KL}}(P\|Q) = H(P, Q) - H(P). \tag{104}$$

#### E. Special Case

If $Q = P$, then

$$D_{\mathrm{KL}}(P\|P) = 0, \tag{105}$$

since $H(P, P) = H(P)$.

#### F. Interpretation

The KL divergence measures the excess cross-entropy relative to the true entropy:

$$D_{\mathrm{KL}}(P\|Q) \geq 0, \tag{106}$$

with equality if and only if $P = Q$ almost everywhere.

### IX. TRPCAGE RESULTS

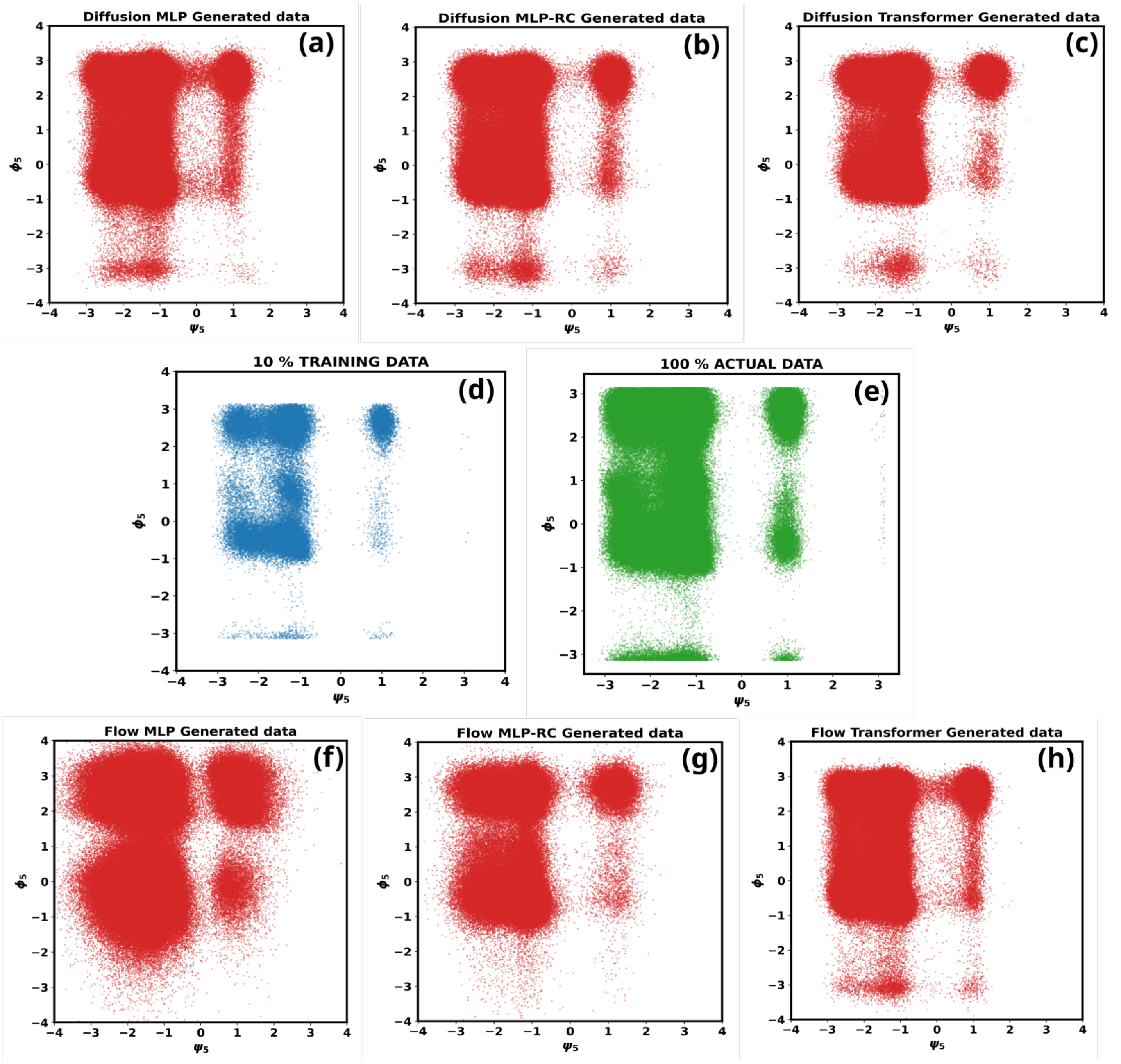


FIG. S1. **Representative** $(\psi_5, \phi_5)$ **marginals for Trp-cage.** Panels (d) and (e) show the training subset and the full reference ensemble. Panels (a)–(c) correspond to diffusion models and panels (f)–(h) to rectified-flow models. Diffusion reproduces the marginal structures with comparatively weak dependence on architecture, whereas RF displays a strong sensitivity to representational capacity and requires Transformer-level expressivity to recover the target pairwise structure reliably.

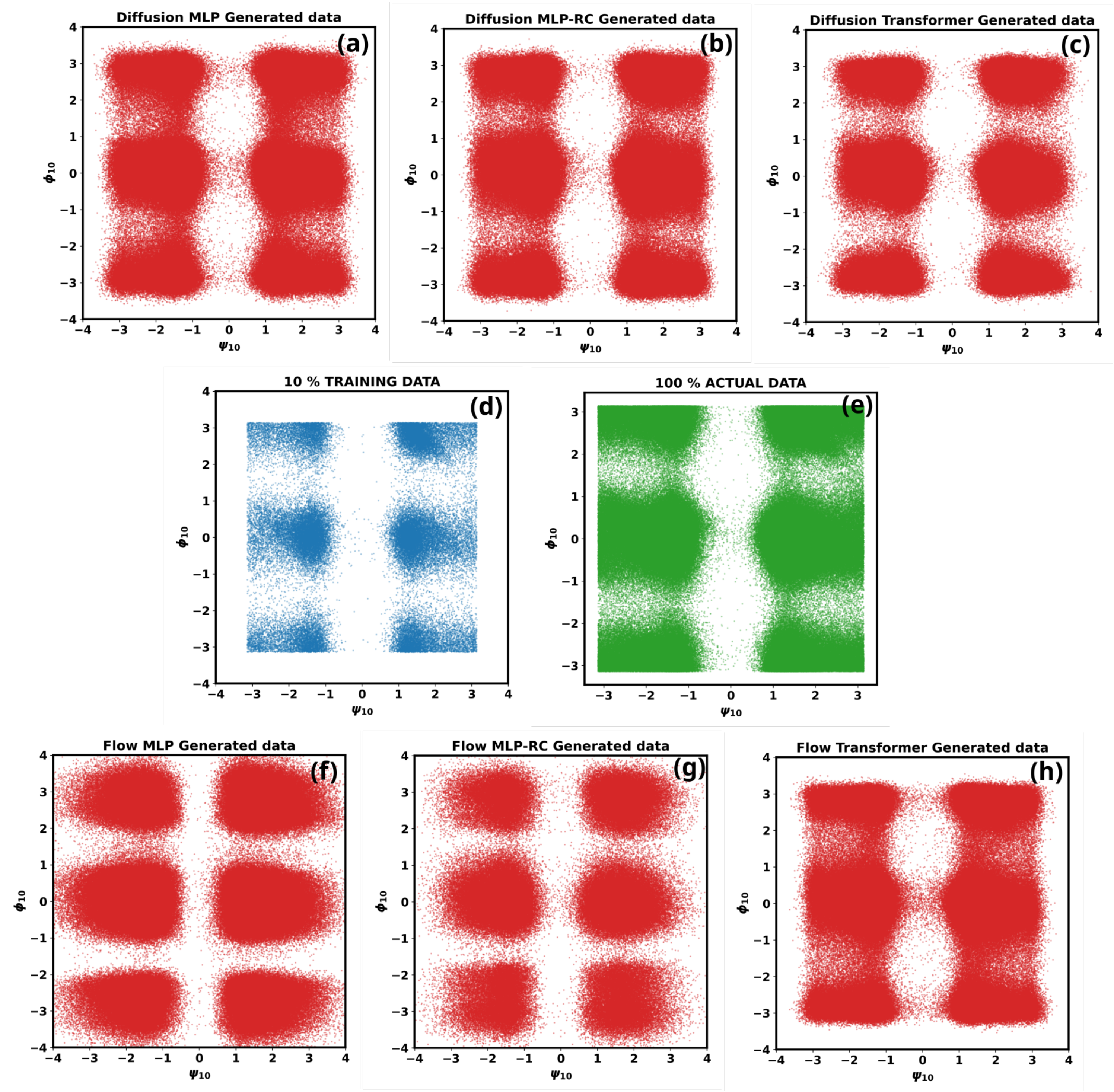


FIG. S2. **Representative** $(\psi_10, \phi_10)$ **marginals for Trp-cage.** Panels (d) and (e) show the training subset and the full reference ensemble. Panels (a)–(c) correspond to diffusion models and panels (f)–(h) to rectified-flow models. Diffusion reproduces the marginal structures with comparatively weak dependence on architecture, whereas RF displays a strong sensitivity to representational capacity and requires Transformer-level expressivity to recover the target pairwise structure reliably.

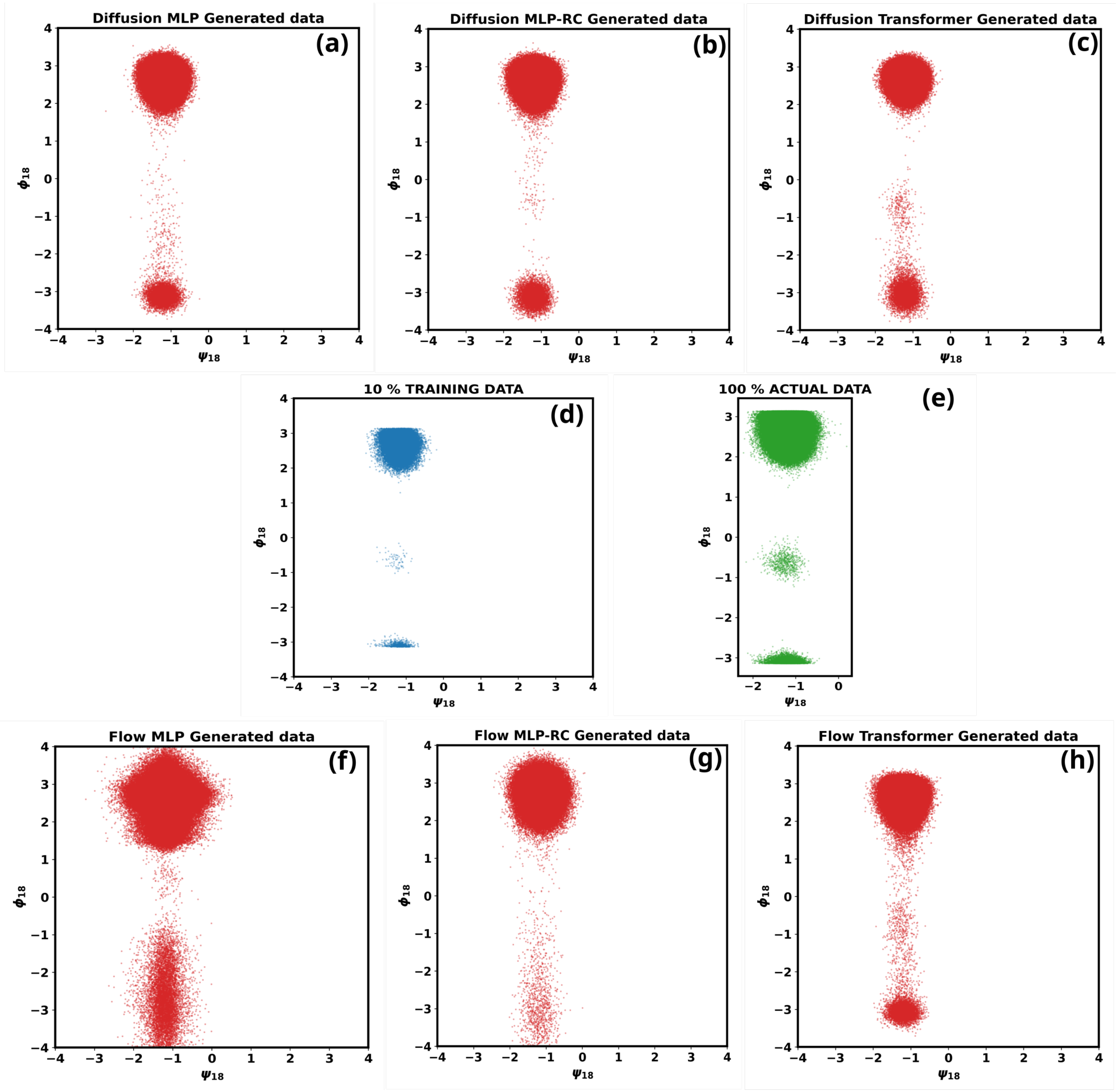


FIG. S3. **Representative** $(\psi_1 8, \phi_1 8)$ **marginals for Trp-cage.** Panels (d) and (e) show the training subset and the full reference ensemble. Panels (a)–(c) correspond to diffusion models and panels (f)–(h) to rectified-flow models. Diffusion reproduces the marginal structures with comparatively weak dependence on architecture, whereas RF displays a strong sensitivity to representational capacity and requires Transformer-level expressivity to recover the target pairwise structure reliably.

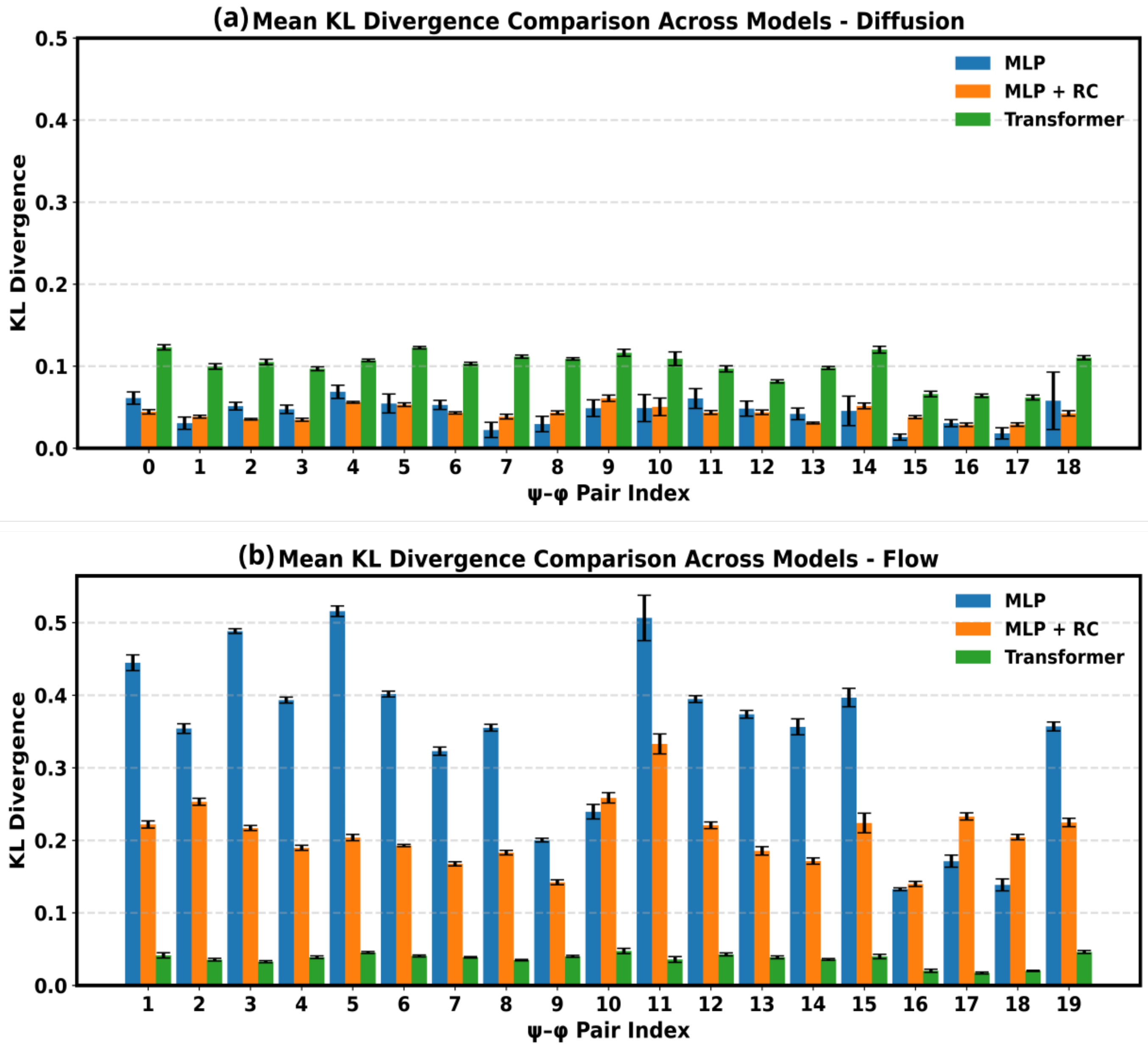


FIG. S4. **Error study of KL divergence across selected Trpcage dihedral pairs using 10% training data using 10 sampled samples . Panel (a) shows diffusion and panel (b) rectified flow. Lower KL indicates better agreement between generated and reference pair distributions. Diffusion remains in a comparatively low-KL regime across architectures, whereas RF exhibits a pronounced capacity bottleneck that is relieved only by the Transformer architecture.Deviations in sampled samples is very less in all architectures across both diffusion and flow models.**

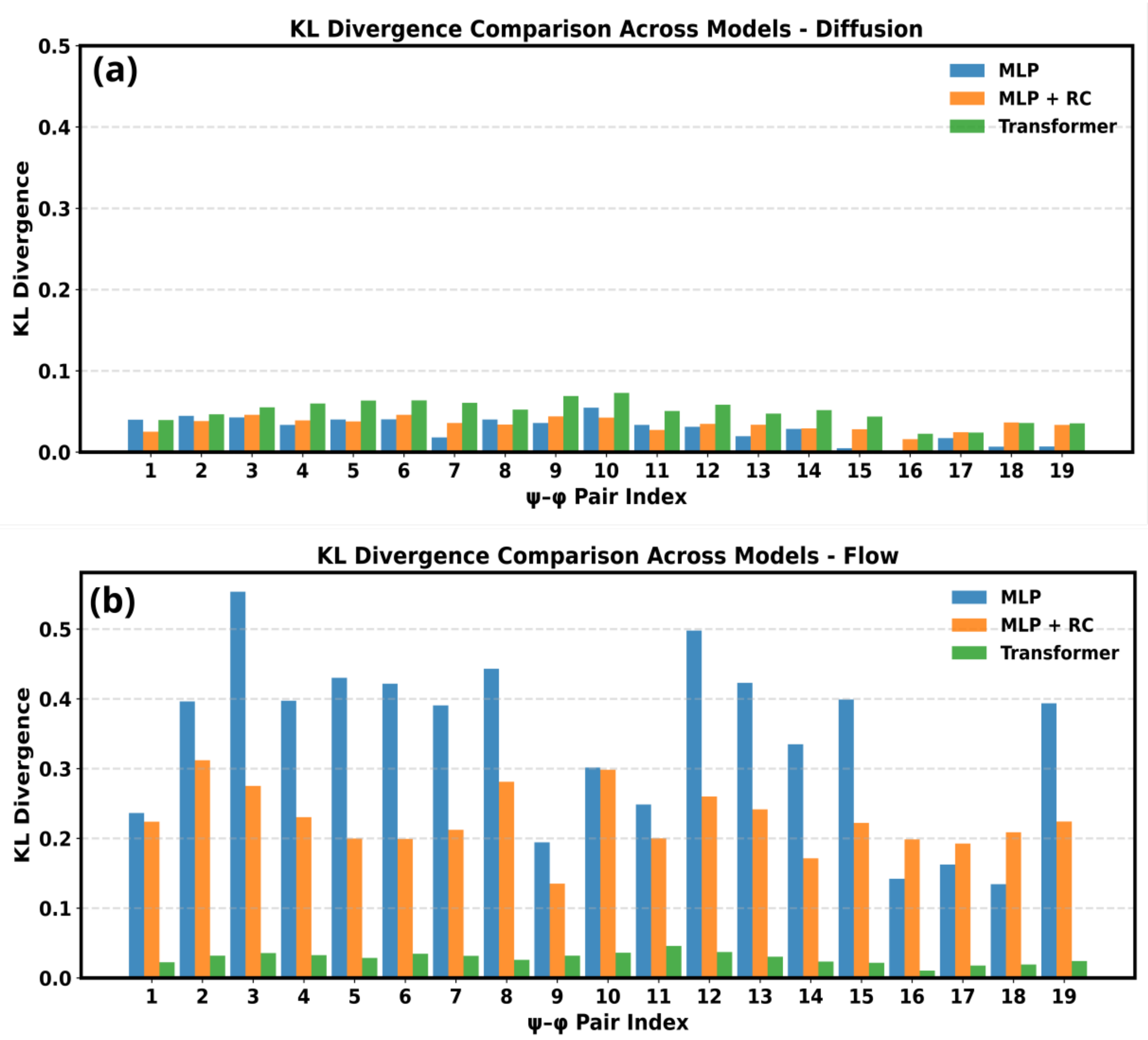


FIG. S5. **KL divergence across selected Trpcage dihedral pairs using 30% training data . Panel (a) shows diffusion and panel (b) rectified flow. Lower KL indicates better agreement between generated and reference pair distributions. Diffusion remains in a comparatively low-KL regime across architectures, whereas RF exhibits a pronounced capacity bottleneck that is relieved only by the Transformer architecture.**

### X. $\alpha$−SYNUCLEIN RESULTS

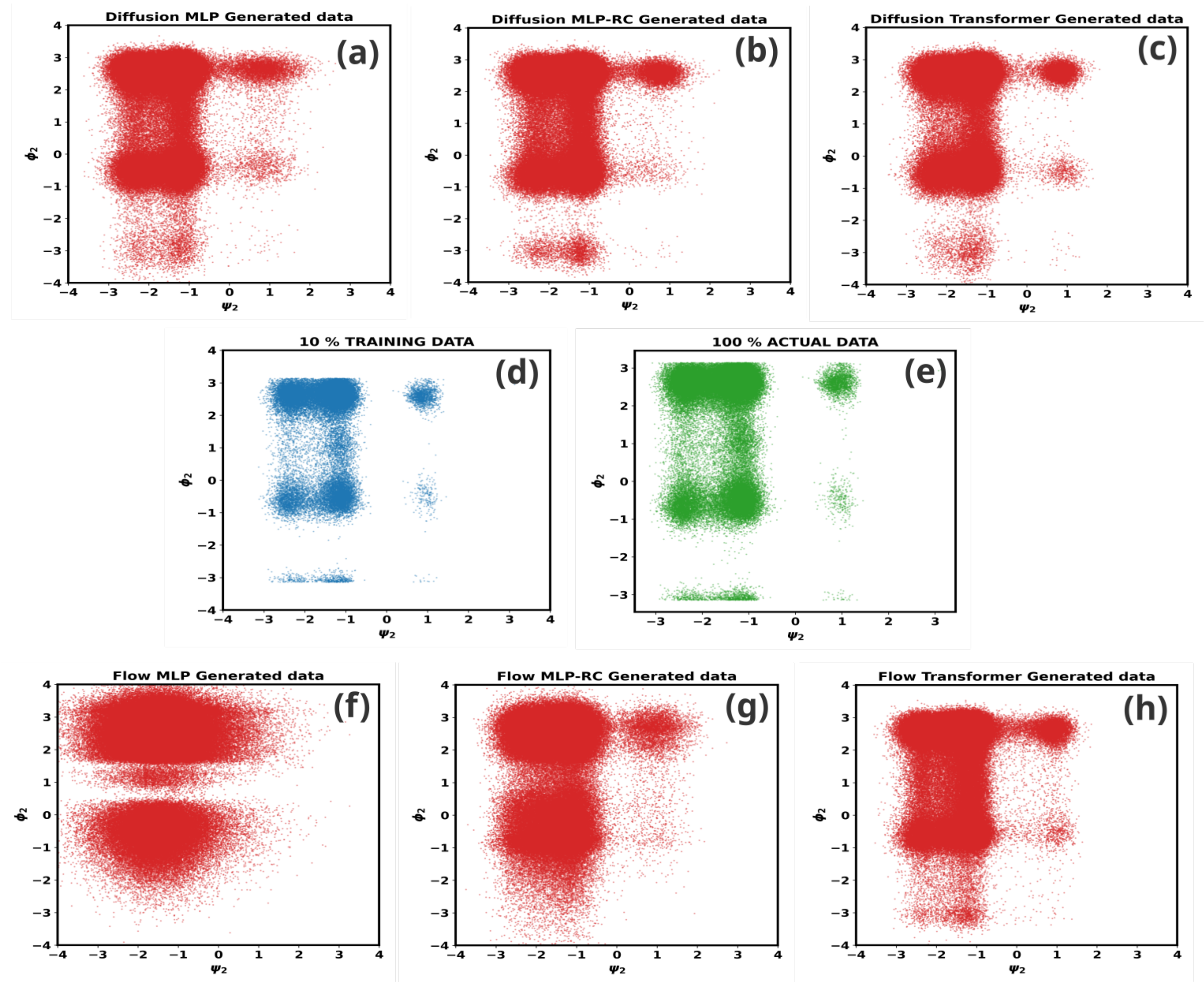


FIG. S6. **Representative $(\psi_2, \phi_2)$ marginals for $\alpha$-synuclein. Panels (d) and (e) show the training subset and the full reference ensemble. Panels (a)–(c) correspond to diffusion models and panels (f)–(h) to rectified-flow models. In this broader and more heterogeneous IDP conformational landscape, diffusion remains relatively robust, whereas RF becomes sharply sensitive to architectural capacity and requires a Transformer to recover the target marginals with reasonable fidelity.**

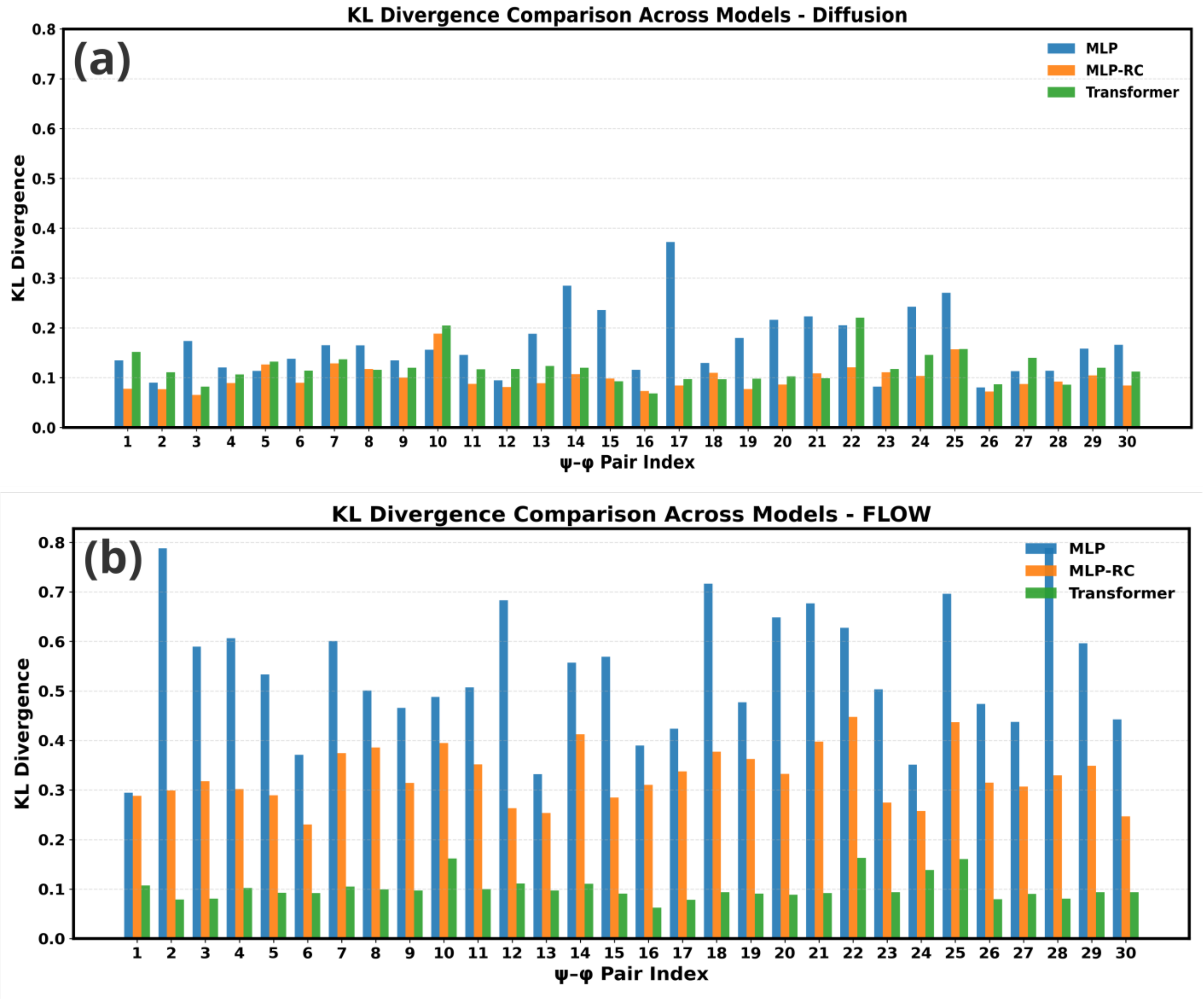


FIG. S7. **KL divergence across selected $\alpha$-synuclein dihedral pairs. Panel (a) shows diffusion and panel (b) rectified flow. The broader spread of KL values reflects the greater heterogeneity of the IDP ensemble. Diffusion remains comparatively stable across architectures, whereas RF exhibits a strong capacity bottleneck that is relieved only by the Transformer architecture.**

### XI. TRPCAGE SAMPLING DYNAMICS

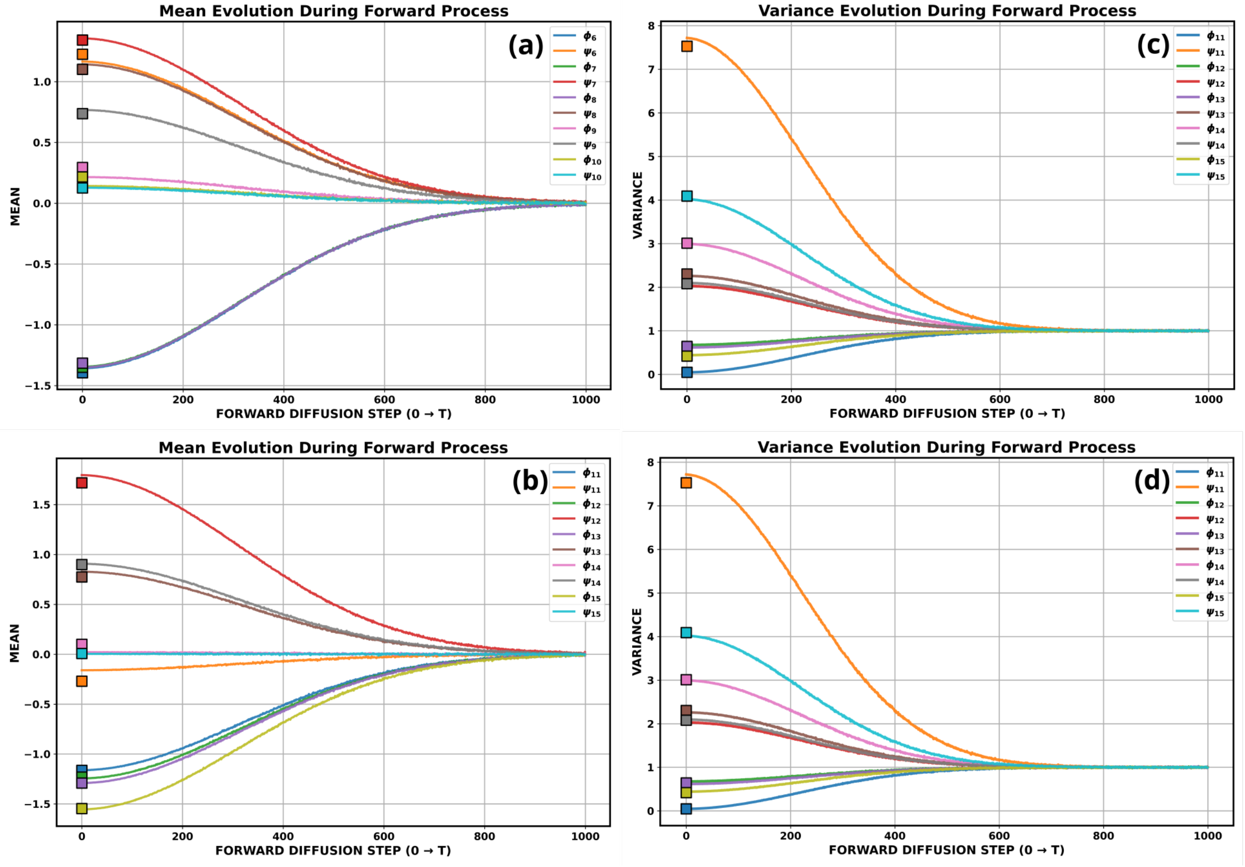


FIG. S8. **Mean and variance convergence to noise during forward diffusion process for Trp-cage** Panels (a) For $(\phi_6, \psi_6, \phi_7, \psi_7, \phi_8, \psi_8, \phi_9, \psi_9, \phi_{10}, \psi_{10})$ and (b) For $(\phi_{11}, \psi_{11}, \phi_{12}, \psi_{12}, \phi_{13}, \psi_{13}, \phi_{14}, \psi_{14}, \phi_{15}, \psi_{15})$

show the evolution of mean values across selected dihedral dimensions, and panels (c) For $(\phi_6, \psi_6, \phi_7, \psi_7, \phi_8, \psi_8, \phi_9, \psi_9, \phi_{10}, \psi_{10})$. and (d) For $(\phi_{11}, \psi_{11}, \phi_{12}, \psi_{12}, \phi_{13}, \psi_{13}, \phi_{14}, \psi_{14}, \phi_{15}, \psi_{15})$

show the corresponding variances. Square markers indicate the reference values obtained from the full MD ensemble. Forward Diffusion brings the mean and variance to the gaussian noise of values 0 and 1 respectively.

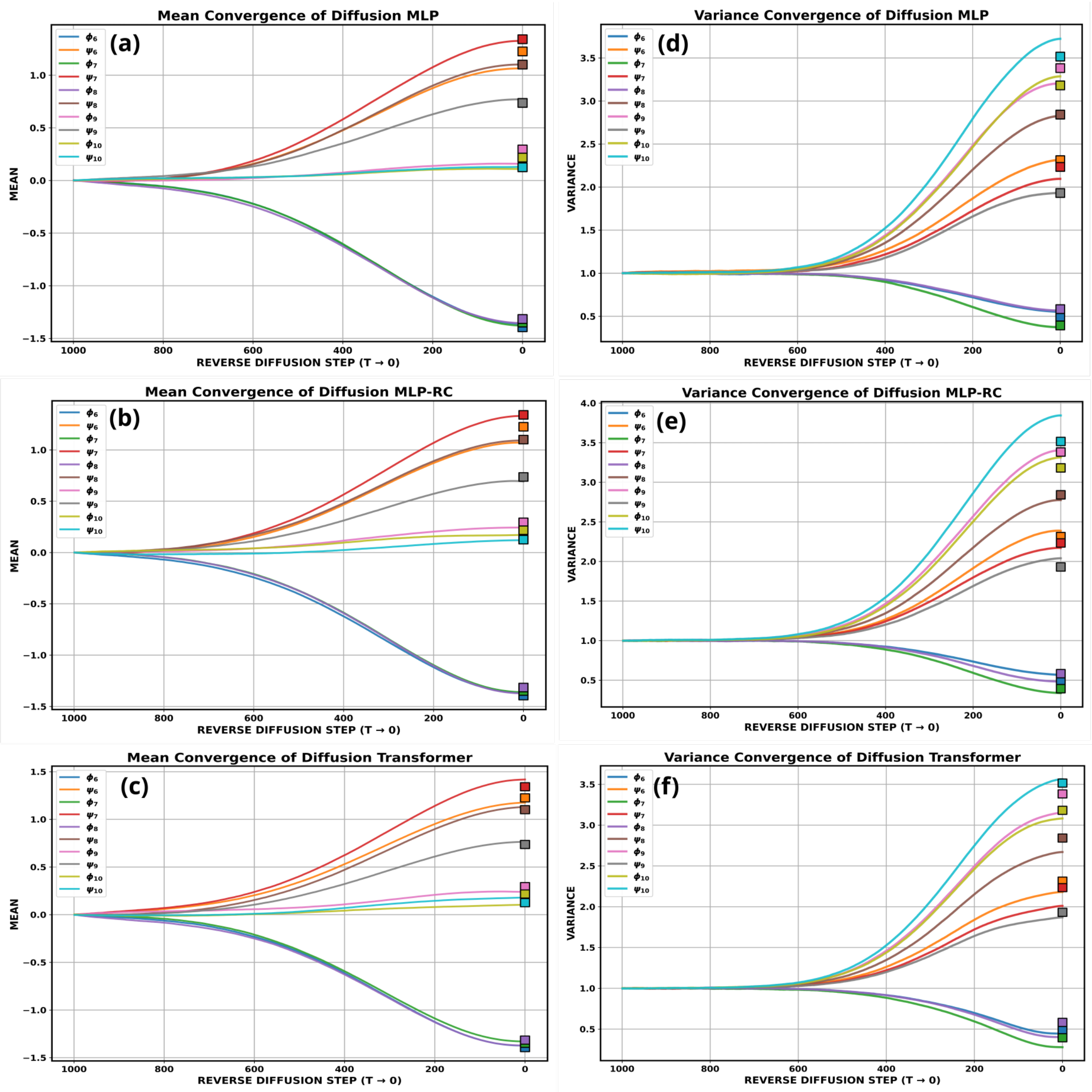


FIG. S9. **Mean and variance convergence during diffusion sampling for Trp-cage of** $(\phi_6, \psi_6, \phi_7, \psi_7, \phi_8, \psi_8, \phi_9, \psi_9, \phi_{10}, \psi_{10})$**.** Panels (a)–(c) show the evolution of mean values across selected dihedral dimensions, and panels (d)–(f) show the corresponding variances. Square markers indicate the reference values obtained from the full MD ensemble. Diffusion brings the means into alignment across all architectures, but variance convergence is most accurate for the residual architecture, indicating more faithful reproduction of conformational fluctuations.

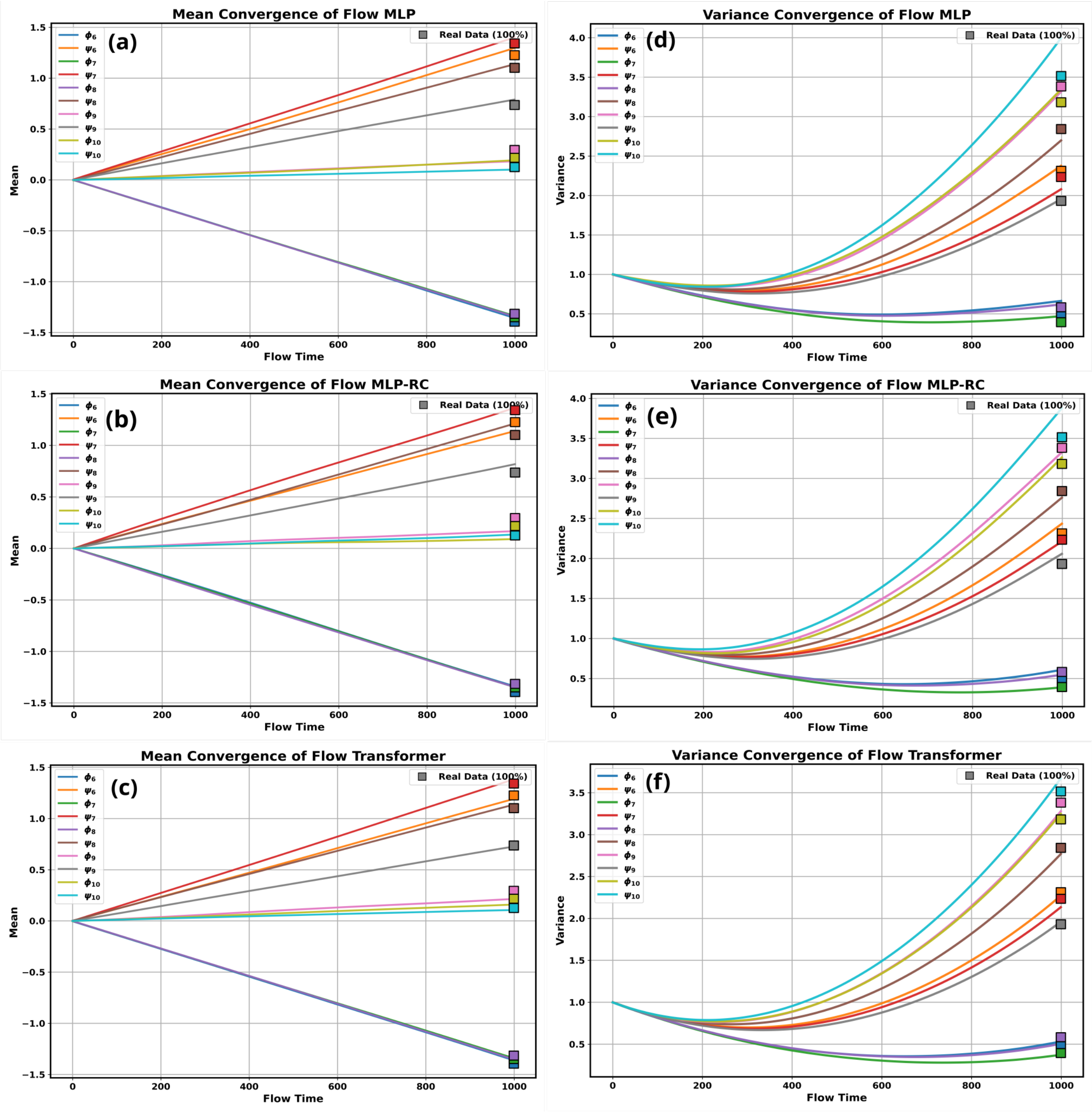


FIG. S10. **Mean and variance convergence during rectified-flow sampling for Trp-cage of** $(\phi_6, \psi_6, \phi_7, \psi_7, \phi_8, \psi_8, \phi_9, \psi_9, \phi_{10}, \psi_{10})$**.** Panels (a)–(c) show mean convergence and panels (d)–(f) show variance convergence. Compared with diffusion, the deterministic transport process is far more sensitive to architecture. The Transformer follows the reference means and variances most closely, while the simpler architectures exhibit persistent deviations, reflecting imperfect transport of both probability location and fluctuation amplitude.

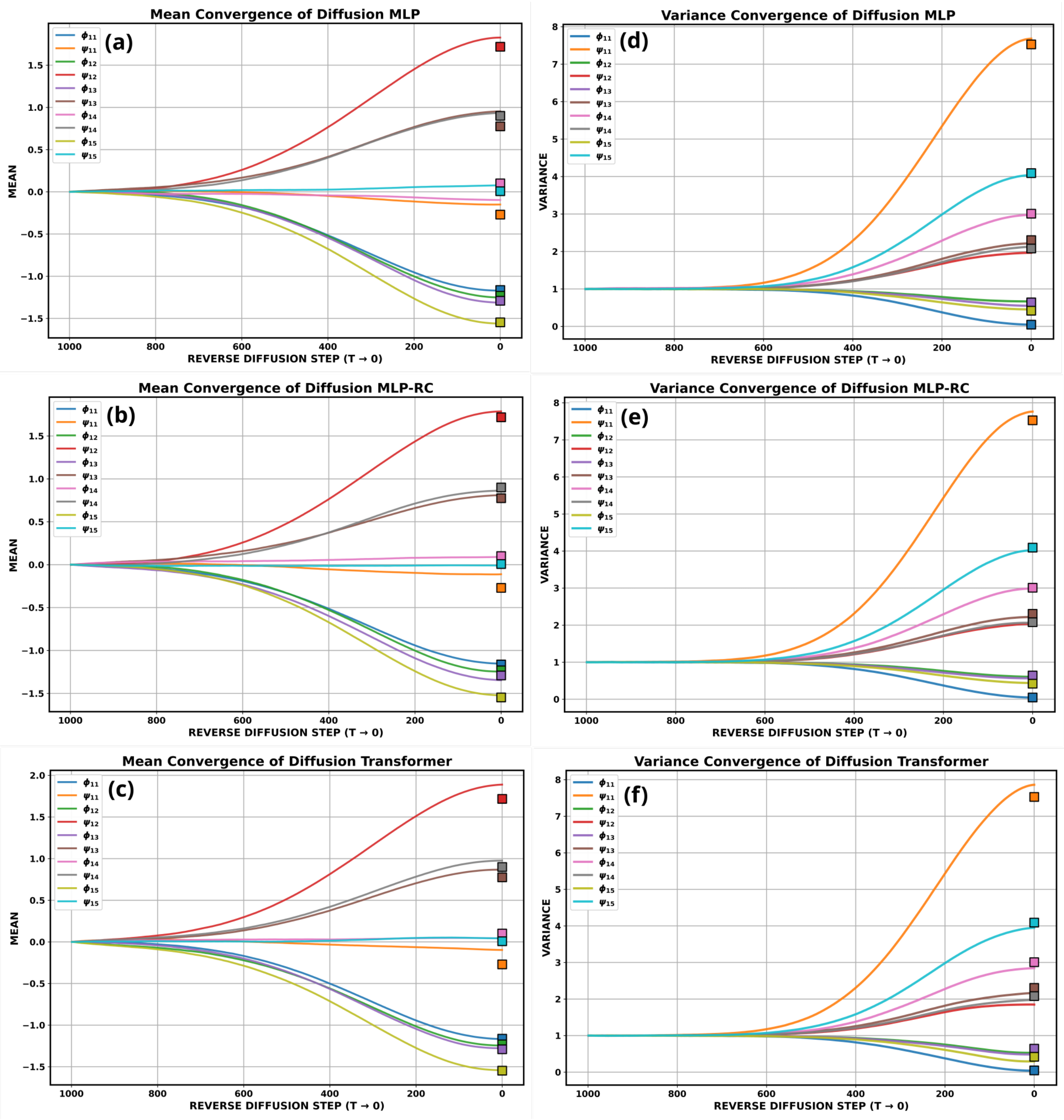


FIG. S11. **Mean and variance convergence during diffusion sampling for Trp-cage of $(\phi_{11}, \psi_{11}, \phi_{12}, \psi_{12}, \phi_{13}, \psi_{13}, \phi_{14}, \psi_{14}, \phi_{15}, \psi_{15})$.** Panels (a)–(c) show the evolution of mean values across selected dihedral dimensions, and panels (d)–(f) show the corresponding variances. Square markers indicate the reference values obtained from the full MD ensemble. Diffusion brings the means into alignment across all architectures, but variance convergence is most accurate for the residual architecture, indicating more faithful reproduction of conformational fluctuations.

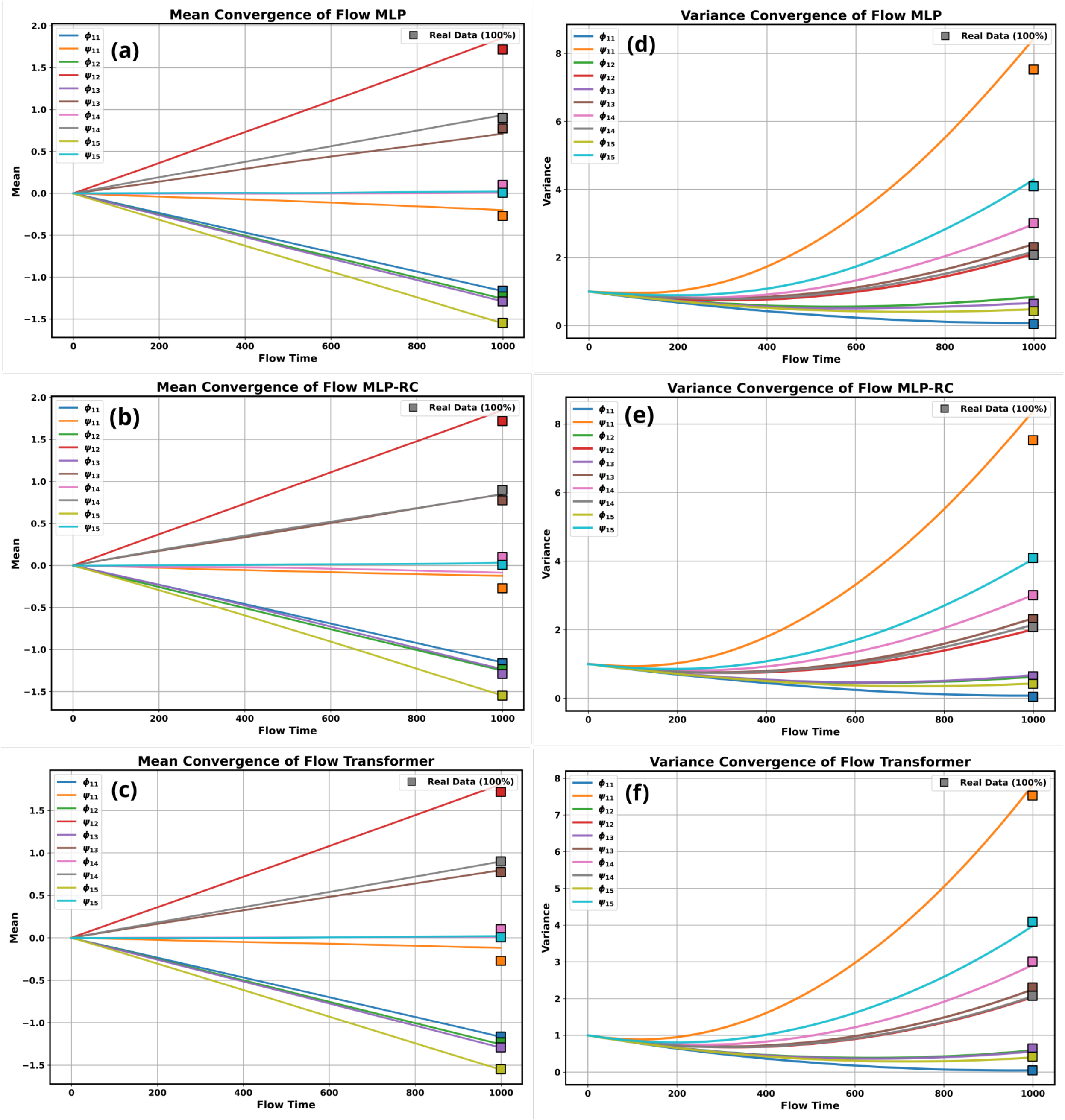


FIG. S12. **Mean and variance convergence during rectified-flow sampling for Trp-cage of** $(\phi_{11}, \psi_{11}, \phi_{12}, \psi_{12}, \phi_{13}, \psi_{13}, \phi_{14}, \psi_{14}, \phi_{15}, \psi_{15})$**.** Panels (a)–(c) show mean convergence and panels (d)–(f) show variance convergence. Compared with diffusion, the deterministic transport process is far more sensitive to architecture. The Transformer follows the reference means and variances most closely, while the simpler architectures exhibit persistent deviations, reflecting imperfect transport of both probability location and fluctuation amplitude.

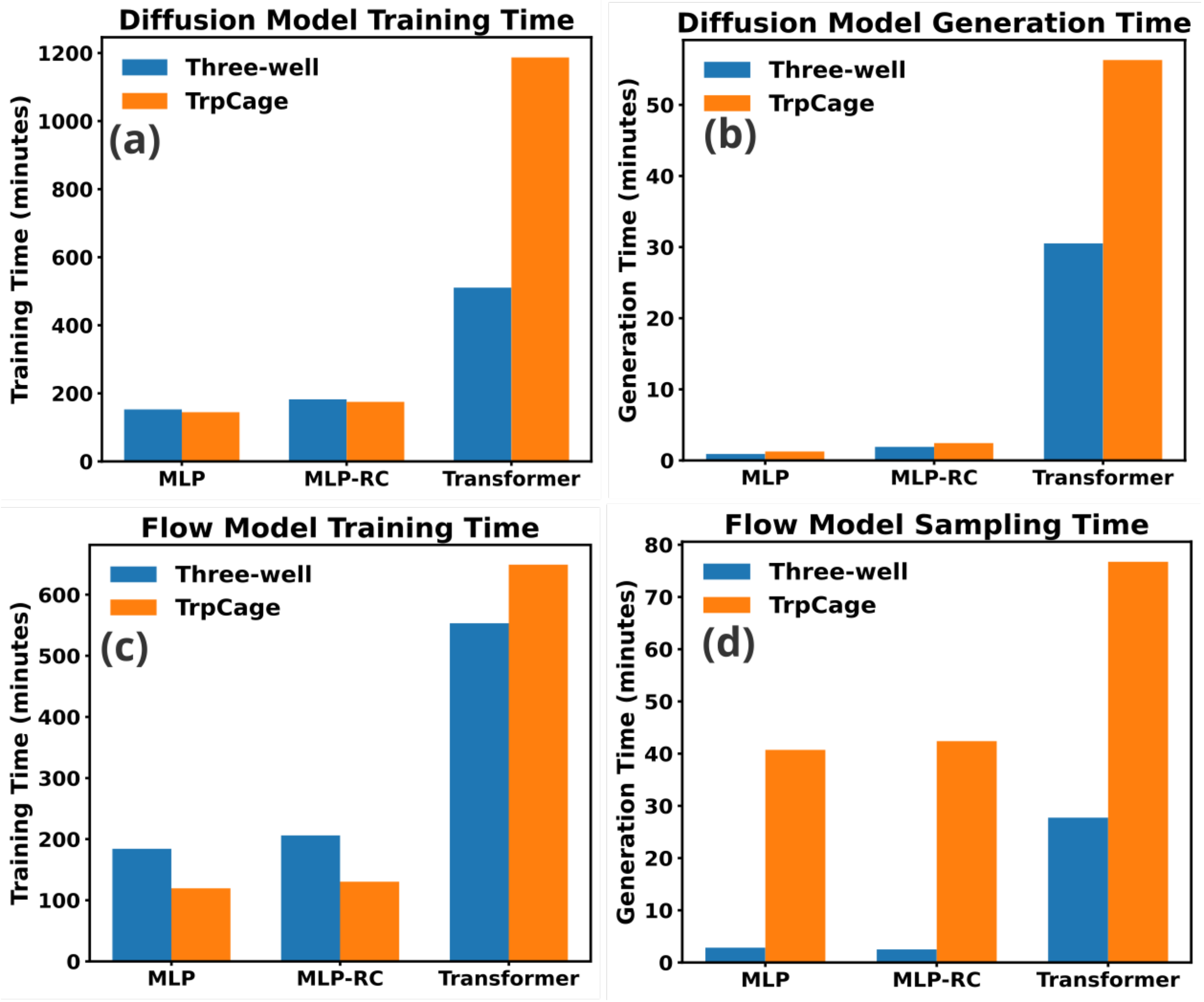


FIG. S13. **Training and sampling time for diffusion and rectified-flow models.** Panels (a) and (b) show the training and sampling times for diffusion, whereas panels (c) and (d) show the corresponding times for rectified flow, for the three-well and Trp-cage systems. In the present implementation, RF is cheaper to train, whereas the Transformer architecture is the most expensive for both paradigms. These trends highlight that practical model selection involves a tradeoff among fidelity, robustness, and computational cost.
